# Whole-body Super-resolution Functional and Molecular Imaging with Panoramic Photoacoustic–Ultrasound Tomography

**DOI:** 10.64898/2026.08.28.747673

**Authors:** Rui Yao, Irma Husain, Jinhuan Luo, Haoming Huo, Xueyi Cai, Nanchao Wang, Tri Vu, Jingting Li, Yirui Xu, Luca Menozzi, Joseph Yang, Matthew Lowerison, Xunrong Luo, Pengfei Song, Junjie Yao

## Abstract

Photoacoustic (PA) and ultrasound (US) imaging provide complementary molecular, functional, and anatomical contrasts. Here, we present a panoramic PA-US imaging platform that integrates multispectral PA computed tomography (PACT) along with reflection-mode and transmission-mode US imaging through a single shared full-ring ultrasound array. We employ an ultrafast planewave transmission scheme in reflection-mode US for power Doppler (PWD) imaging and ultrasound localization microscopy (ULM). Additionally, we use the transmission-mode US to reconstruct a spatially resolved speed of sound (SoS) map that corrects both PA and US reconstruction. Such correction sharpens the resolution of PACT, suppresses the artifacts of PWD, and improves microbubble localization of ULM. Elevational scanning further enables whole-body volumetric imaging with co-registered PA and US contrasts. The integrated system maps photoswitchable DrBphP1-expressing tumors alongside their blood perfusion and oxygenation environment. Applying the platform to monitor unilateral renal ischemia– reperfusion injury, we report that microvascular perfusion and renal oxygenation recover at different rates. Collectively, we demonstrate that the integrated PA-US imaging platform provides a unified framework for multiparametric study of anatomy, perfusion, microvascular flow, oxygenation, and molecular activities.

## Introduction

Photoacoustic computed tomography (PACT) and ultrasound (US) imaging are closely related imaging modalities that provide complementary views of biological tissues. In PACT, short laser pulses are delivered into tissue, where the optical absorption by endogenous or exogenous chromophores induces transient thermoelastic expansion and generates broadband acoustic waves [1, 2]. By detecting the emitted acoustic waves with an array of ultrasonic transducers and solving an acoustic inverse problem, PACT maps the spatial distribution of optical absorption deep within scattering tissue. In contrast, US pulse-echo imaging actively transmits acoustic pulses and detects the echoes generated by spatial variations in tissue’s acoustic impedance, thereby providing structural information based on acoustic scattering. Despite their distinct contrast mechanisms, PACT and US share many core hardware and signal processing components [3, 4]: both rely on broadband acoustic detection, radiofrequency data acquisition, acoustic propagation models, and similar image reconstruction methods. In both modalities, image quality is strongly affected by the detector bandwidth, angular coverage, acoustic attenuation, and the assumed speed of sound (SoS) used during reconstruction [5–8].

Because of these shared acoustic foundations and complementary contrast mechanisms, integrating PACT and US into a single imaging platform is a natural and powerful strategy [3, 4, 9–22]. From an instrumentation perspective, although various optical acoustic sensors with high detection sensitivity and broad bandwidth have been developed for PACT [23–26], they lack the acoustic transmission capability. By contrast, piezoelectric transducer arrays and data acquisition electronics can generally be readily shared between PACT and US. From a biological perspective, PACT provides rich optical absorption contrast and is particularly sensitive to blood oxygenation and molecular probes, whereas US provides label-free anatomical imaging, contrast-enhanced vascular imaging, and quantitative blood flow measurements. In particular, US B-mode can delineate tissue morphology, microbubble (MB)-enhanced power Doppler (PWD) imaging can visualize blood perfusion with high contrast, and ultrasound localization microscopy (ULM) can resolve microvascular structures beyond the acoustic diffraction limit by localizing and tracking individual MBs across successive frames [27]. Integrating these capabilities enables co-registered assessment of tissue anatomy, molecular composition, oxygenation, vascular architecture, and blood flow. Such multiparametric imaging is particularly valuable for preclinical studies of cancer, vascular biology, organ injury, metabolism, and therapeutic response, in which disease progression is rarely captured by a single contrast mechanism alone [3].

The performance of both PACT and US, however, depends critically on detector geometry [6, 28]. Linear-array transducers are widely available and highly compatible with conventional US imaging, but their limited angular aperture results in limited-view artifacts [29] and anisotropic spatial resolution. These limitations are particularly pronounced in PACT, in which acoustic waves emitted by targets that are poorly oriented relative to the detection aperture are weakly detected or entirely missed. Matrix and hemispherical arrays improve angular sampling and enable volumetric imaging, but they can be costly and provide a limited field of view (FOV) [30, 31]. Ring-array transducers provide an attractive alternative for cross-sectional small-animal imaging because they offer a large FOV and full 360° in-plane acoustic detection. This full-view geometry improves angular sampling, reduces limited-view artifacts, and provides more uniform in-plane resolution and sensitivity than the other one-sided detection geometries [28, 32–34]. Enclosed apertures have also been tested for multimodal operation [15–17, 22] and, in addition to conventional PACT and US pulse-echo imaging, can enable transmission-mode ultrasound tomography (UST) across the object to estimate SoS and acoustic-attenuation distributions. Incorporating the measured SoS distribution into image reconstruction is important for reducing phase errors [17, 35, 36]. These errors are especially consequential for fine vascular structures, causing blurring, spatial distortion, or even duplication of the same structure. Thus, ring-array geometry not only provides improved acoustic angular coverage but also provides a framework for more accurate reconstruction of both PA and US images.

Combining PACT with US, the ring-array detection geometry and SoS estimation have been explored before—for example, in transmission–reflection optoacoustic ultrasound of whole mice [15, 16] and in ring-array PACT/US with adaptive or dual-speed reconstruction [17, 22, 34]. Our group and others have also integrated PACT with ULM for brain and tumor imaging [9, 12, 14, 20]. Here, the same full-ring array that performs multispectral molecular PACT also measures the object’s SoS field by UST and feeds that measured field back into both PACT and US reconstruction, rather than assuming a homogeneous or two-speed medium. To our knowledge, this is the first demonstration of transmission-measured, spatially resolved SoS correction applied to *in vivo* MB-enhanced PWD and super-resolution ULM, co-registered with PA molecular contrast.

Here, we present an integrated PA-US imaging platform based on a full-ring transducer array for whole-body imaging of small animals (**Fig. 1a**). The system combines multispectral PACT, US B-mode, MB-enhanced PWD, and ULM within a unified acquisition framework, with transmission-mode UST providing spatially resolved SoS maps for acoustic-heterogeneity correction. Additionally, we use elevational scanning to extend the imaging FOV and enable volumetric visualization of anatomical structures, vascular functions, and molecular activities. We demonstrate the capabilities of the system in three representative applications: whole-body neck-to-tail imaging of a mouse *in vivo*, functional and molecular imaging of tumor xenografts, and longitudinal assessment of renal ischemia-reperfusion injury (IRI). In the IRI model, the multiparametric capability reveals that ULM-measured microvascular perfusion and PA- measured oxygenation recover on different timescales after injury—a functional divergence that no single contrast mechanism would capture.

**Figure 1.**
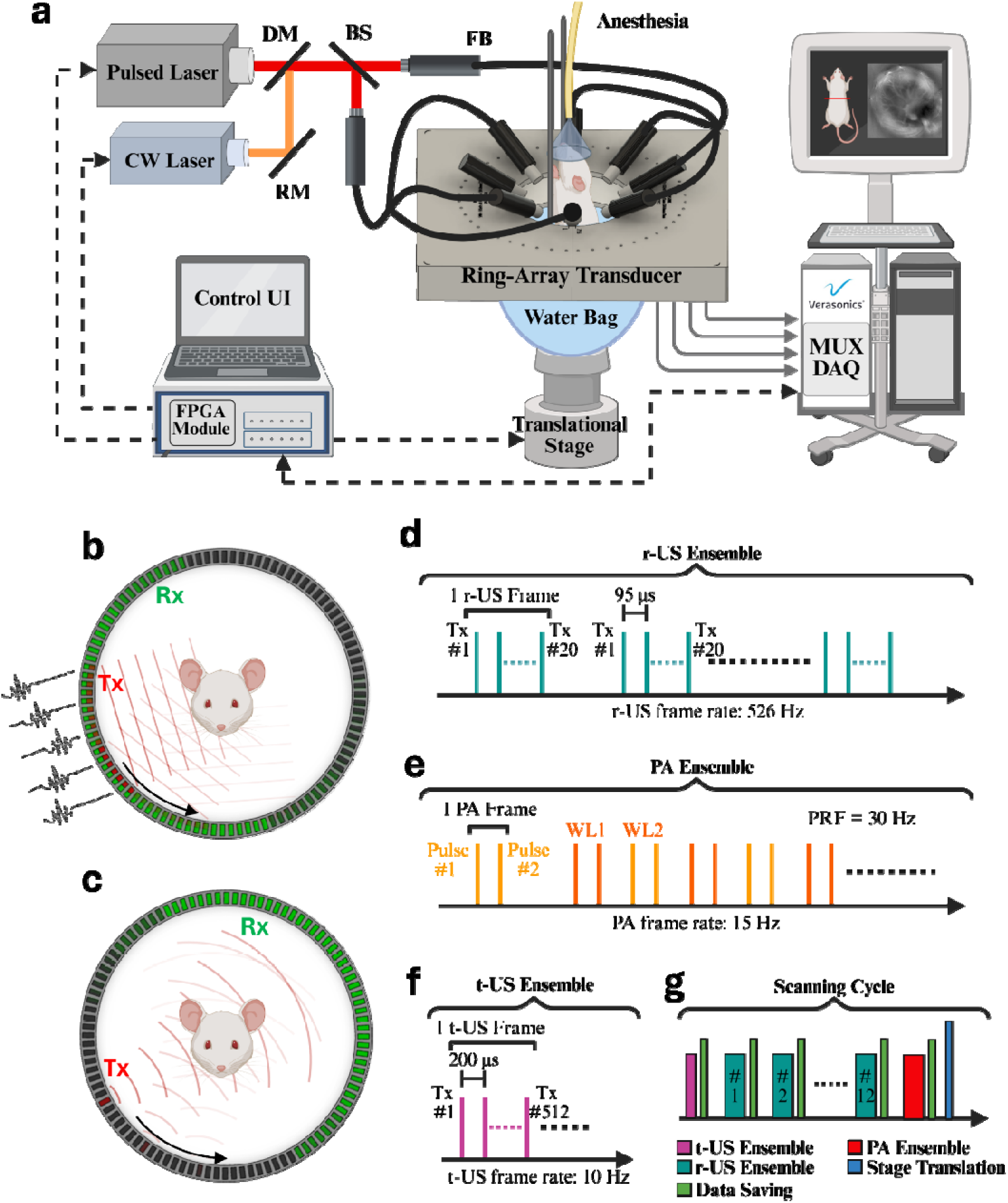
Full-ring PA-US imaging system and multimodal data-acquisition sequence. **(a)** Schematic of the imaging system. BS, beam splitter; CW, continuous-wave laser; DAQ, data-acquisition system; DM, dichroic mirror; FB, fiber bundle; MUX, multiplexer; RM, reflective mirror. **(b)** Plane-wave transmission and receiving scheme for reflection-mode ultrasound (r-US). The transmission (Tx) aperture is shown in red, and the receiving (Rx) aperture is shown in green. Element-specific transmission delays were applied across the Tx aperture to generate a plane wave. **(c)** Single-element transmission and opposite-side receiving scheme for transmission-mode ultrasound (t- US). **(d)** Structure of an r-US slow-time ensemble. Each ensemble contained 250 compounded frames, and each frame comprised 20 plane-wave Tx/Rx events, yielding a compounded frame rate of 526 Hz. **(e)** Structure of a PA ensemble. The number of PA frames varied among experiments. Because the 512-element array was multiplexed to a 256-channel DAQ, each full-ring PA frame required two consecutive acquisitions at the same wavelength, resulting in an effective frame rate of 15 Hz. **(f)** Structure of a t-US ensemble. Each ensemble contained three frames, and each frame comprised 512 single-element Tx/Rx events distributed around the full-ring aperture. **(g)** Data-acquisition sequence for one scanning cycle. At each elevational position, t-US, r-US, and PA ensembles were acquired sequentially. After acquisition and data transfer were completed, the translational stage advanced to the next scanning position.

## Results

### Integrated PA-US Imaging Using the Ring-Array System

The imaging system was built around a 512-element full-ring ultrasonic transducer array with a center frequency of 5 MHz (**Fig. 1a**). The transducer exhibited a one-way receive bandwidth of >100% and a two-way transmit–receive bandwidth of approximately 60% (**Supplementary Fig. 1**) and was described in our earlier study [37]. US acquisition comprised two modes: reflection-mode US (r-US) and transmission-mode US (t-US). For r-US, we employed a plane-wave transmission scheme consisting of 20 consecutive transmissions distributed around the full-ring aperture (**Fig. 1b** and **Supplementary Fig. 2a**). In each transmit/receive (Tx/Rx) event, the Tx and Rx apertures were located on the same side of the ring and shared the same center point. Coherent compounding of the 20 plane-wave transmissions [38] yielded a B-mode frame rate of 526 Hz, enabling the tracking of rapidly moving MBs required for ULM [39]. In contrast, t-US acquisition used 512 single-element transmissions, with the Tx and Rx apertures positioned on opposite sides of the ring (**Fig. 1c** and **Supplementary Fig. 2c**). Representative simulated pressure fields for r-US and t-US transmission are shown in **Supplementary Fig. 3**. For PA imaging, optical excitation was delivered through circumferentially arranged fiber bundles coupled to a pulsed optical parametric oscillator (OPO) laser (**Fig. 1a**). A 635-nm continuous-wave (CW) laser was used only for photoswitching experiments. Because the 512-element array was multiplexed to a 256-channel data acquisition system, acquisition of one full-ring PA frame required two laser pulses, each corresponding to signals received by 256 elements. Consequently, the PA frame rate was 15 Hz, half the 30-Hz pulse repetition frequency (PRF) of the OPO laser (see **Methods**). At each scanning position, t-US, r-US, and PA ensembles were acquired sequentially (**Figs. 1d–1f**), after which the translational stage advanced to the next position (**Fig. 1g**). 3D datasets were generated by stacking the reconstructed 2D cross-sectional images acquired at successive elevational positions.

The spatial resolution of the system was characterized by imaging two orthogonally oriented copper wires with a nominal diameter of 10 µm. Cross-sectional images of two selected regions of interest (ROIs; **Supplementary Figs. 4a–4c** and **5a–5c**) showed that PA and US B-mode imaging provided similar in-plane radial resolutions. Based on the full width at half maximum (FWHM) of the signal envelope profiles, the radial resolution ranged from 0.25 to 0.29 mm and remained relatively uniform between the two ROIs (**Supplementary Figs. 4d, 4f, 5d**, and **5f**). The elevational resolution of B-mode ranged from 0.88 to 1.01 mm (**Supplementary Figs. 4e** and **4g**), whereas that of PA ranged from 1.16 to 1.17 mm (**Supplementary Figs. 5e** and **5g**). The moderately finer elevational resolution of B-mode imaging is consistent with two-way elevational focusing during transmission and receiving, compared with receiving-only elevational focusing in PACT. Furthermore, because of the transducer’s relatively small elevational aperture, the elevational resolution was approximately four times worse than the in-plane radial resolution.

### PA and r-US Image Reconstruction Assisted by t-US-derived SoS Map

We incorporated t-US measurements to estimate the spatial SoS distribution and compensate for acoustic heterogeneity during PA and r-US image reconstruction (**Figs. 2a**–**2c**; see **Methods** for details). Representative cross-sectional PA images acquired at the excitation wavelength of 1064 nm and PWD images of the mouse abdominal region were reconstructed using either a single homogeneous SoS of 1521 m/s in a 36 °C water bath, or a spatially varying SoS map derived from t-US measurements (**Fig. 2**). We additionally compared SoS-map-assisted reconstruction with a dual-SoS approach [17, 34], in which different homogeneous SoS values were assigned to the animal body and surrounding water (**Supplementary Fig. 7**). The SoS distributions used for these reconstruction approaches are shown in **Supplementary Fig. 6**.

**Figure 2.**
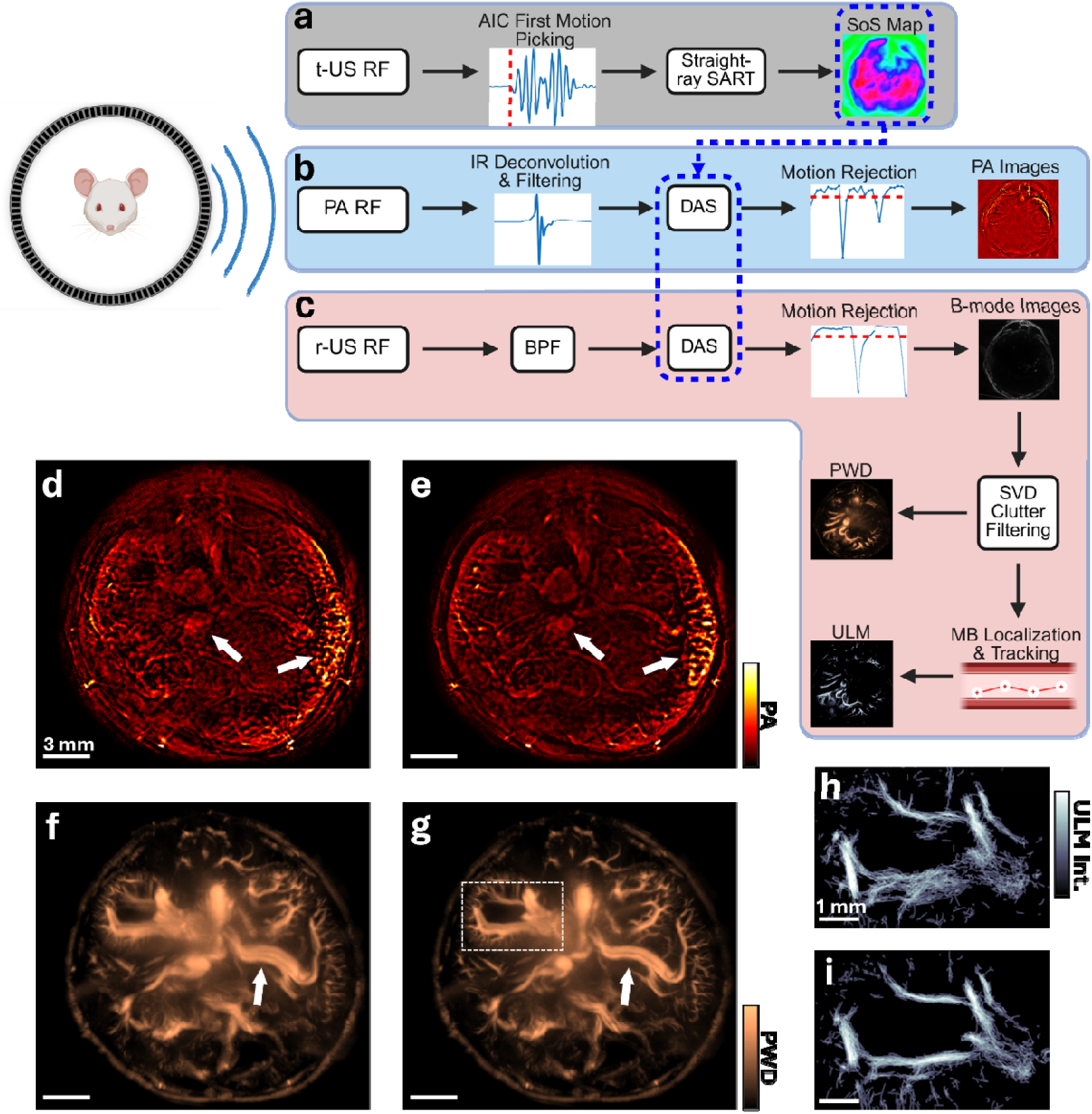
SoS-corrected PA-US image reconstruction framework. Image reconstruction and processing workflows for **(a)** transmission-mode ultrasound (t-US), **(b)** photoacoustic (PA) imaging, and **(c)** reflection-mode ultrasound (r-US). Acquired signals were recorded as radiofrequency (RF) data. The t-US measurements were used to reconstruct a spatial speed-of-sound (SoS) map, which was incorporated into delay- and-sum (DAS) beamforming for both PA and r-US reconstruction to correct acoustic propagation-time errors. The r-US data were further processed to generate B-mode, power Doppler (PWD), and ultrasound localization microscopy (ULM) images. SART, simultaneous algebraic reconstruction technique; AIC, Akaike information criterion; IR, impulse response; BPF, bandpass filter; SVD, singular value decomposition. Representative cross-sectional **(d, e)** PA images acquired at 1064 nm and **(f, g)** PWD images reconstructed using **(d, f)** a single homogeneous SoS of 1521 m/s, or **(e, g)** a spatially varying t-US-derived SoS map. The homogeneous value of 1521 m/s corresponded to the SoS of the water bath at 36 °C. **(h, i)** Magnified ULM images of the boxed right-kidney region in panel **(g)**, reconstructed using **(h)** a single homogeneous SoS of 1521 m/s or **(i)** the t-US-derived SoS map. SoS-map-assisted reconstruction improved PA, PWD, and ULM images.

In the PA images, vascular signals near the skin surface and within deeper tissue were more sharply focused when the t-US-derived SoS map was incorporated into reconstruction (**Fig. 2e**) than when a single homogeneous SoS was assumed (**Fig. 2d**). Features such as the portal vein and spleen, indicated by the white arrows in **Fig. 2d**, appeared blurred under the single-SoS assumption but were better resolved after SoS correction. Dual-SoS reconstruction also provided effective phase-error compensation (**Supplementary Fig. 7a**). However, because this approach assigned a uniform SoS of 1550 m/s to the entire animal body, it produced poorer focusing than the spatially varying SoS-map-assisted reconstruction in selected regions (**Supplementary Fig. 7e**).

The improvement in PWD image quality was most apparent for deeper vessels. For example, the splenic artery and splenic vein, indicated by the white arrows in **Figs. 2f** and **2g**, appeared as three separate vascular features under the single-SoS assumption (**Fig. 2f**), indicating duplication caused by propagation-time errors. This artifact was substantially reduced after reconstruction with the t-US-derived SoS map (**Fig. 2g**). Dual-SoS reconstruction produced results comparable to those obtained with the spatially varying SoS map (**Supplementary Fig. 7c**), although the latter provided better vessel separation in selected regions (**Supplementary Fig. 7f**).

The benefits of SoS correction were also evident in the ULM images. In the right-kidney region, SoS-map-assisted reconstruction more clearly resolved the major renal vessels and cortical microvasculature (**Fig. 2i**), whereas these features were less distinct under the single-SoS assumption (**Fig. 2h**). The ULM image sharpness was also increased after SoS correction as a result of improved MB localization and tracking.

### Whole-Body Neck-to-Tail Imaging *in vivo*

To demonstrate the volumetric imaging capability of the ring-array platform, we performed neck-to-tail imaging of the mouse torso using 200 scanning cycles over a total elevational range of 6 cm (**Fig. 3**). During *in vivo* imaging, the anesthetized mouse was positioned vertically at the center of the ring array in a temperature-controlled water bath maintained at 36 °C (**Fig. 1a**). MBs (Definity) were continuously infused through the tail vein to maintain a relatively stable circulating MB concentration throughout the imaging session.

**Figure 3.**
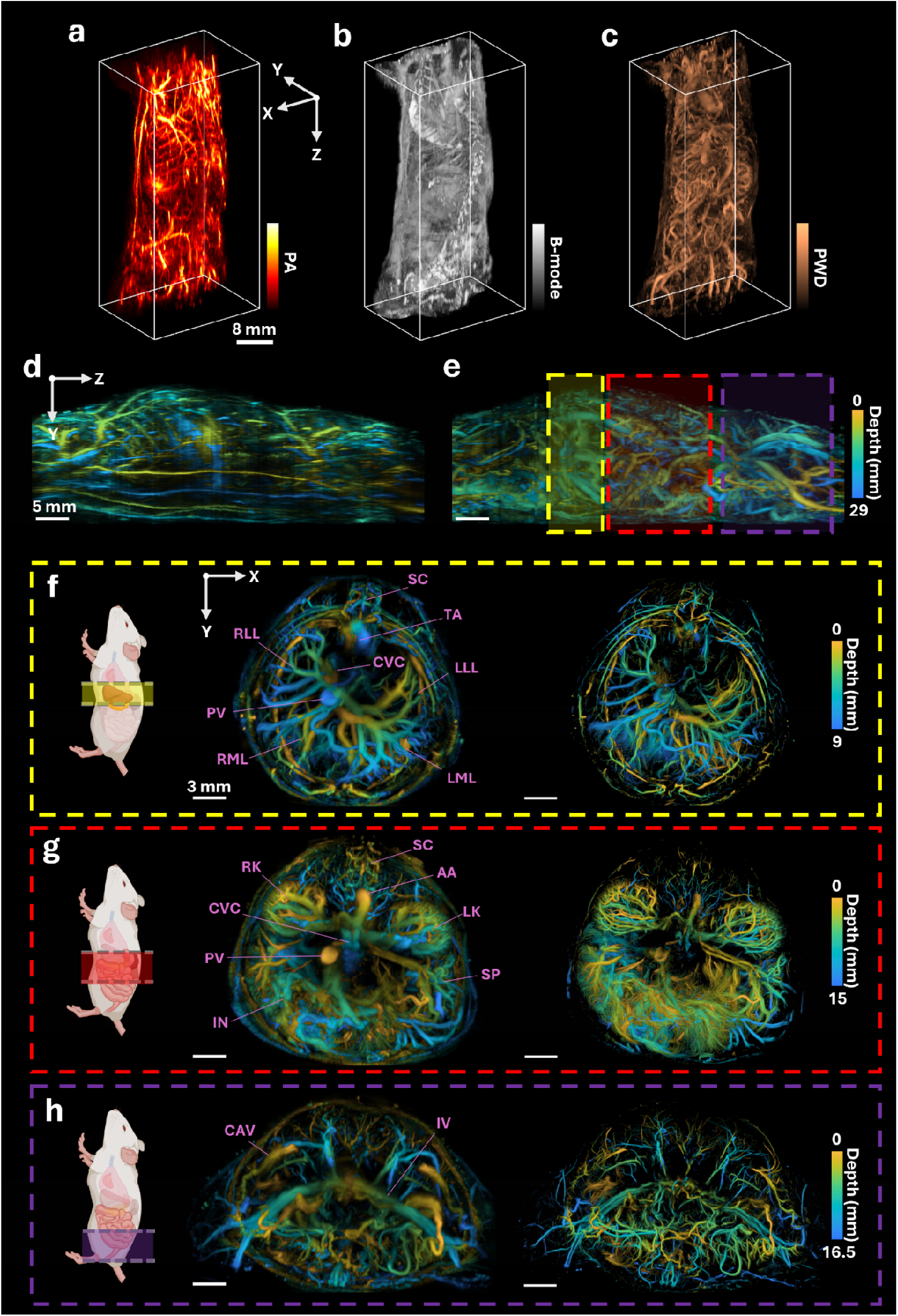
*In vivo* neck-to-tail PA-US imaging of the mouse torso. 3D volumetric renderings of **(a)** PA imaging acquired at 870 nm, **(b)** US B-mode, and **(c)** MB-enhanced PWD. **(d, e)** Depth-encoded sagittal MAPs of the **(d)** PA and **(e)** PWD volumes, showing vascular structures from the thoracic cavity to the lower abdomen. Color represents the depth along the viewing axis. **(f–h)** Depth-encoded transverse PWD (left) and ULM MAPs (right), corresponding to the color-coded regions indicated in panel **(e)**: **(f)** upper abdomen, **(g)** mid-abdomen, and **(h)** lower abdomen. The ULM images provide improved delineation of microvascular structures relative to PWD images. SC, spinal cord; TA, thoracic aorta; CVC, caudal vena cava; RLL, right lateral lobe of the liver; RML, right medial lobe of the liver; LML, left medial lobe of the liver; LLL, left lateral lobe of the liver; PV, portal vein; AA, abdominal aorta; RK, right kidney; LK, left kidney; SP, spleen; IN, intestines; CAV, cranial abdominal vein; IV, iliac vessels.

Representative 3D renderings are shown in **Figs. 3a**–**3c** and **Supplementary Videos 1-3**. PACT images at 870 nm primarily highlighted blood-rich structures, with greater sensitivity to oxygenated hemoglobin at this wavelength (**Fig. 3a** and **Supplementary Video 1**) [40]. US B-mode images delineated the external body boundary and internal anatomical structures like the vertebral column based on acoustic-scattering contrast (**Fig. 3b** and **Supplementary Video 2**). MB-enhanced PWD imaging visualized perfused vasculature throughout the imaging volume, extending from the thoracic cavity to the lower abdominal region (**Fig. 3c** and **Supplementary Video 3**). Depth-encoded sagittal maximum amplitude projections (MAPs) further illustrate the neck-to-tail vascular distributions captured by PACT (**Fig. 3d**) and PWD (**Fig. 3e**).

The left column of **Figs. 3f**-**3h** shows transverse PWD MAPs from three representative regions. In the upper abdomen, vessels associated with four liver lobes—the right lateral lobe (RLL), right medial lobe (RML), left medial lobe (LML), and left lateral lobe (LLL)—were clearly visualized (**Fig. 3f**). In the mid-abdomen, PWD images depicted vasculature associated with the right and left kidneys, intestines, and spleen (**Fig. 3g**). In the lower abdomen, the iliac and other lower-abdominal vessels were visualized (**Fig. 3h**). Additional sliding-depth MAPs from neck to tail for PACT, US B-mode, and PWD were shown in **Supplementary Videos 4-6**, respectively. Because the elevational resolution was approximately four times the in-plane radial resolution, small vessels were generally more clearly visible in the transverse images than in the sagittal images.

The corresponding ULM images in the right column of **Figs. 3f**–**3h** provided a more detailed depiction of the microvascular architecture. Improvements in vessel delineation relative to PWD imaging were evaluated in two ROIs encompassing the left kidney and iliac vessels (**Supplementary Fig. 8**). Compared with PWD imaging, ULM produced sharper vessel profiles and better separation of close vessels, allowing the iliac artery, iliac vein, and small renal interlobar vessels to be distinguished. However, because the volumetric datasets were assembled from independently acquired cross-sectional images, the MB tracking algorithm could not reliably track MBs traveling predominantly along the elevational (or along the spine) direction. Consequently, although most in-plane vasculature was clearly depicted, vessels oriented primarily along the elevational direction, including the aorta, caudal vena cava, and portal vein, were not captured well in the ULM images (**Figs. 3f**–**3h**).

### Functional and Molecular PA-US Imaging of Tumor Xenografts

To demonstrate molecular imaging beyond endogenous hemoglobin contrast, we performed PA photoswitching imaging in a mouse bearing bilateral 4T1 tumor xenografts. DrBphP1 is a bacterial phytochrome photoreceptor that can be reversibly switched between ON and OFF states through wavelength-dependent changes in its absorption spectrum [41]. In this study, DrBphP1-expressing 4T1 cells (DrBphP1-4T1) were implanted into one mammary fat pad of the mice, whereas wild-type 4T1 cells (WT-4T1) were implanted contralaterally as a negative control (**Fig. 4a**). *In vivo* fluorescence imaging confirmed DrBphP1 expression in the DrBphP1-4T1 tumor, whereas the WT-4T1 tumor exhibited minimal fluorescence signal (**Fig. 4b**).

**Figure 4.**
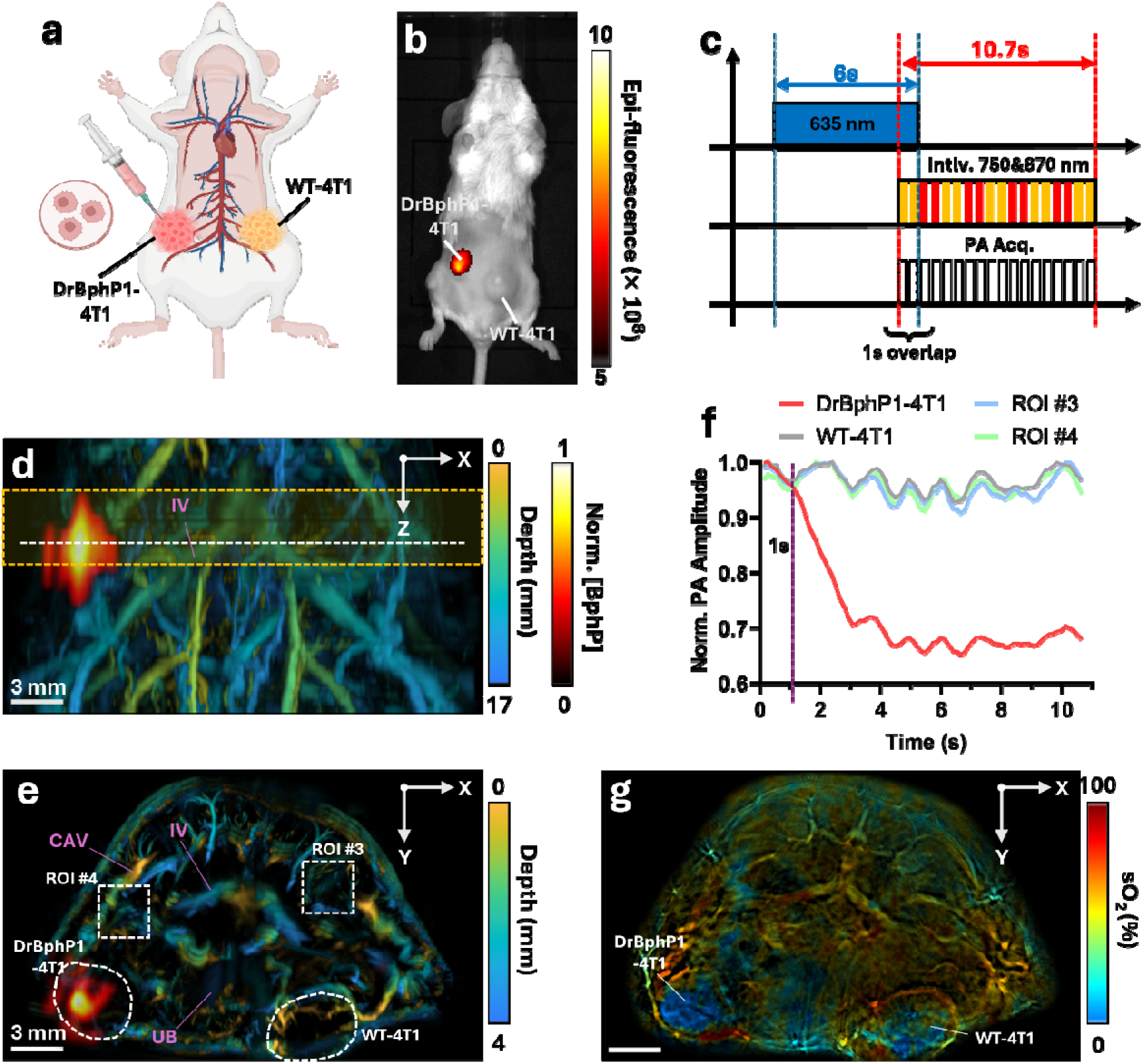
***In vivo* functional and photoswitchable molecular imaging of a tumor-bearing mouse. (a)** Schematic of bilateral orthotopic implantation of DrBphP1- expressing 4T1 cells and wild-type 4T1 (WT-4T1) cells into the mammary fat pads near the iliac vessels (IV). **(b)** Representative *in vivo* IVIS fluorescence image showing strong DrBphP1-associated fluorescence in the DrBphP1-4T1 tumor and no fluorescence in the contralateral WT-4T1 tumor. **(c)** Illumination and PA acquisition sequence for imaging the DrBphP1 photoswitching response. The mouse was illuminated with 635- nm continuous-wave light for 6 s to switch DrBphP1 to the ON state. Interleaved PA acquisition at 750 and 870 nm began 5 s after the onset of 635-nm illumination, resulting in a 1-s overlap, and continued for approximately 10.7 s. **(d)** Coronal PWD MAP overlaid with the isolated DrBphP1-specific differential PA signals. **(e)** Transverse MAP of the yellow-boxed region in panel **(d)**, showing the two tumors and surrounding vasculature. CAV, cranial abdominal vein; IV, iliac vessels; UB, urinary bladder. **(f)** Normalized PA signal amplitudes from the four ROIs indicated in panel **(e)** over the data acquisition. The vertical dashed line indicates the end of 635-nm illumination. Only the DrBphP1-4T1 tumor exhibited a pronounced decrease in PA signal amplitude following the photoswitching illumination. **(g)** PA cross-section image color-coded by estimated blood oxygen saturation (sO₂), showing decreased oxygenation within the central regions of the two tumors. The elevational position of the cross-section is indicated by the white dashed line in panel **(d)**.

For photoswitching, 635-nm CW light was used to switch DrBphP1 to its ON state, followed by 750-nm pulsed illumination to transition it to the OFF state (**Fig. 4c**). PA signals were acquired at interleaved wavelengths of 750 and 870 nm. The 750-nm measurements captured the photoswitching response of DrBphP1, whereas the combined 750- and 870-nm measurements were used to estimate blood oxygenation (sO₂). Because endogenous absorbers (e.g., hemoglobin, lipids, collagen) were not expected to exhibit the same signal switching response to the illumination sequence, DrBphP1-specific contrast was isolated by subtracting the OFF-state PA image from the ON-state image (see **Methods**). The resulting differential PA image was overlaid on the PWD image to show the spatial relationship between DrBphP1 expression and the surrounding vasculature. In the coronal MAP, strong differential PA contrast was detected in the DrBphP1-4T1 tumor near the iliac vessels but not in the contralateral WT-4T1 tumor (**Fig. 4d**). A transverse MAP provided an additional view of the two tumors surrounding the urinary bladder and showed reduced perfused-vessel density within their central regions (**Fig. 4e** and **Supplementary Video 7**). The tumor boundaries were also delineated in the corresponding US B-mode cross-section (**Supplementary Fig. 9, Supplementary Video 8**). A sO₂-encoded PA cross-section image further showed lower blood oxygenation within the central regions of both tumors (**Fig. 4g**). To confirm the temporal photoswitching response, we compared the PA signal amplitudes from the two tumors and two additional control ROIs indicated in **Fig. 4e**. The PA signal from the DrBphP1-4T1 tumor was switched to the OFF level within approximately 2 s (**Fig. 4f**). In contrast, the other three ROIs, including the WT-4T1 tumor, exhibited no substantial signal decrease. The synchronous fluctuations observed across all four ROIs were likely associated with respiratory motion (**Supplementary Video 9**).

### Longitudinal Imaging of Renal Ischemia-Reperfusion Injury

To demonstrate the ability of the integrated PA-US platform to monitor organ-specific vascular and functional changes over time, we performed longitudinal imaging in a unilateral renal ischemia-reperfusion injury (IRI) model (**Fig. 5**). Renal IRI is a common cause of acute kidney injury and frequently occurs during kidney transplantation [42]. In this study, blood flow to the right kidney was occluded for 30 min and then restored. Three mice were imaged at four time points: baseline, Day 1 approximately 1.5 h after IRI surgery, Day 7, and Day 13. At each time point, both the injured right kidney and the intact left kidney were evaluated on their vessel density and blood flow by ULM, and sO₂ by PACT.

**Figure 5.**
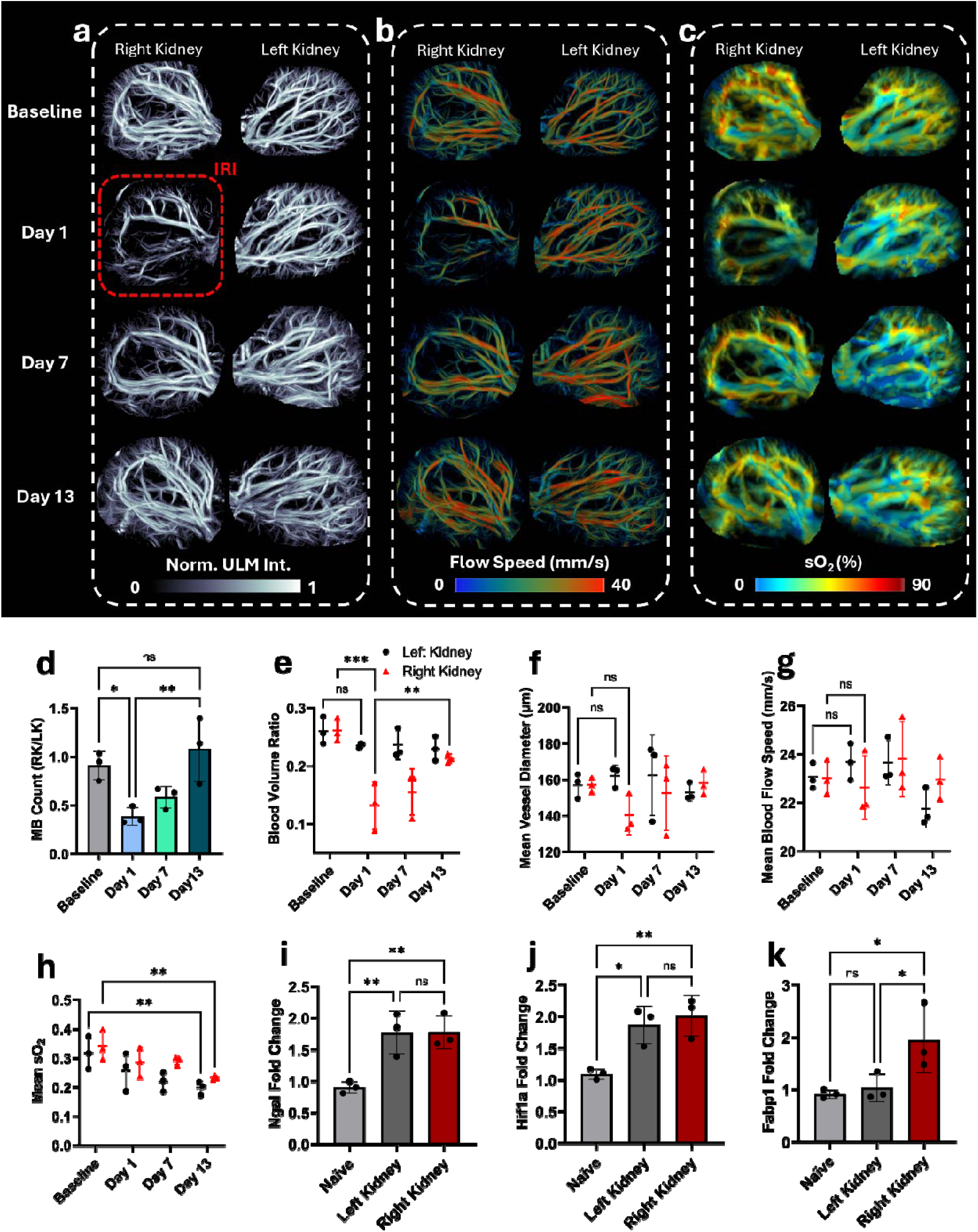
Longitudinal imaging of unilateral renal ischemia–reperfusion injury. Representative **(a)** ULM intensity maps, **(b)** flow-speed-encoded ULM maps, and **(c)**PWD vascular maps color-coded by PA-derived sO₂ for the right and left kidneys at baseline and on Day 1, 7, and 13. The right renal pedicle was occluded for 30 min after baseline imaging, and Day 1 imaging was performed approximately 1.5 h after the IRI procedure. **(d)** Right-to-left kidney MB count ratio. The right-to-left kidney MB count ratio decreased significantly from baseline to Day 1 (p = 0.0279) and increased from Day 1 to Day 13 (p = 0.0086); no significant difference was detected between baseline and Day 13 (p = 0.3760). **(e)** Blood volume ratio, defined as the fraction of the kidney ROI occupied by perfused vessels. The right-kidney blood volume ratio decreased from baseline to Day 1 (p = 0.0002) and increased from Day 1 to Day 13 (p = 0.0059). No significant change in left-kidney blood volume ratio was detected between baseline and Day 1 (p = 0.3344). **(f)** Mean whole-kidney vessel diameter. Mean vessel diameter did not change significantly from baseline to Day 1 in either the left (p = 0.5857) or right (p = 0.0885) kidney. **(g)** Mean whole-kidney blood flow speed. Similarly, mean blood flow speed did not change significantly from baseline to Day 1 in either the left (p = 0.5054) or right (p = 0.6826) kidney. **(h)** Mean whole-kidney sO₂. Mean sO₂ decreased from baseline to Day 13 in both the left (p = 0.0012) and right (p = 0.0026) kidneys. Relative renal expressions of **(i)** *Ngal*, **(j)** *Hif1a*, and **(k)** *Fabp1* measured by RT-qPCR 2 h after unilateral right-kidney IRI. Kidneys from naïve mice served as controls; “left kidney” and “right kidney” denote the contralateral and injured kidneys, respectively, from mice subjected to IRI. *Ngal* expression was higher in both the contralateral left kidney (p = 0.0062) and injured right kidney (p = 0.0060) than in naïve kidneys, with no significant difference between the two kidneys. *Hif1a* expression was also higher in the left (p = 0.0104) and right (p = 0.0048) kidneys than in naïve kidneys, with no significant difference between the two kidneys. *Fabp1* expression was higher in the injured right kidney than in both naïve kidneys (p = 0.0190) and the contralateral left kidney (p = 0.0311); no significant difference was detected between naïve and left kidneys. Data are presented as *mean* ± SD(n = 3). The MB count ratio was analyzed using one-way repeated-measures ANOVA; Blood volume ratio, vessel diameter, blood flow speed, and mean sO₂ were analyzed using two-way repeated-measures ANOVA; *Ngal*, *Hif1a*, and *Fabp1* were assessed using unpaired Student’s t-test. Statistical analyses were performed using GraphPad Prism. *p<0.05; ** p<0.01; ns, not significant.

Representative ULM intensity images from the same mouse showed a marked reduction in MB signals in the injured right kidney on Day 1, indicating an acute decrease in renal perfusion after IRI (**Fig. 5a**). The vascular signal partially recovered by Day 7 and further improved by Day 13. In contrast, vascular structures in the contralateral left kidney remained relatively preserved throughout the experiment period. ULM flow-speed maps showed no clear longitudinal trend in MB flow speed in either kidney across the four time points (**Fig. 5b**). Finally, sO_2_-encoded PWD images showed a slight decrease in overall blood oxygenation level in both kidneys after baseline (**Fig. 5c**).

Quantitative analysis supported these imaging observations. The right-to-left kidney MB count ratio decreased significantly from baseline to Day 1, increased significantly from Day 1 to Day 13, and did not differ significantly between baseline and Day 13 (**Fig. 5d**). This ratio served as a relative indicator of renal perfusion between the two kidneys. Using a relative metric reduced the influence of systemic MB concentration and motion artifacts. A similar temporal pattern was observed in the right-kidney blood volume ratio, defined as the volume occupied by perfused vessels relative to the whole-kidney ROI volume (**Fig. 5e**). The right-kidney blood volume ratio decreased significantly from baseline to Day 1 and subsequently increased by Day 13, whereas the left-kidney value did not differ significantly between baseline and Day 1. Nonetheless, neither kidney showed a significant change in the mean whole-kidney vessel diameter or blood flow speed between baseline and Day 1, and neither metric exhibited a clear longitudinal trend (**Figs. 5f** and **5g**). In contrast, the mean whole-kidney sO₂ was significantly lower at Day 13 than at baseline in both kidneys (**Fig. 5h**). Thus, the quick recovery of perfusion-related ULM metrics in the injured kidney was not accompanied by the recovery in the whole-kidney oxygenation over the observation period. Because sO₂ was estimated from two wavelengths without fluence correction (**Methods**), these oxygenation measurements were interpreted relative to each kidney’s own baseline. The temporal divergence between blood perfusion and oxygenation was therefore reported as a relative trend rather than an absolute measurement.

To provide independent biological validation for renal injury and hypoxia-associated responses, we assessed gene expressions in a separate cohort of mice undergoing the same procedures. These animals were euthanized 2 h after the IRI procedure, and both kidneys were collected for analysis. Kidneys from naïve mice that did not undergo IRI served as controls. We first measured the expression of Neutrophil Gelatinase-Associated Lipocalin (*Ngal)*, an early kidney injury marker produced by kidney tubular cells after hypoxic injury [43, 44]. As expected, *Ngal* expression was significantly elevated in both kidneys of mice subjected to unilateral IRI, compared with kidneys from naïve mice (**Fig. 5i**). Second, we examined the expression of Hypoxia Inducible Factor 1a (*Hif1a*), an oxygen-sensing regulator involved in cellular responses to hypoxia [45]. Similar to *Ngal*, *Hif1a* expression was also significantly elevated in both kidneys after unilateral IRI (**Fig. 5j**). Although the left kidney was not directly subjected to ischemia, the elevated *Ngal* and *Hif1a* expression in the contralateral kidney was consistent with a systemic or compensatory response to unilateral renal injury [46, 47]. Finally, we examined the expression of another injury marker, Fatty Acid Binding Protein 1 (*Fabp1*). *Fabp1* was significantly higher in the injured right kidney than in the contralateral left kidney and kidneys from naïve mice (**Fig. 5k**), supporting its association with renal injury in this model [48].

## Discussion

In this study, we developed an integrated PA-US imaging platform that combines PACT with r-US and t-US within a unified acquisition and reconstruction framework (**Figs. 1 and 2**). By using the same 512-element ring array for all acoustic measurements, the system provides intrinsically co-registered anatomical, vascular, hemodynamic, oxygenation, and molecular information. The full 360° aperture offers broad angular coverage and a large cross-sectional FOV, while elevational scanning extends the imaging range for volumetric imaging. We demonstrated these capabilities through whole-body small animal imaging (**Fig. 3**), photoswitchable molecular and functional imaging of tumor xenografts (**Fig. 4**), and longitudinal monitoring of renal IRI (**Fig. 5**). Collectively, these experiments show that the integrated PA-US platform can characterize biological processes across spatial and functional dimensions that would be difficult to capture using any individual modality alone.

An important feature of the system is the use of t-US to estimate the spatial SoS distribution and correct acoustic propagation-time errors during both PA and r-US reconstruction. This correction was particularly beneficial for vascular imaging, in which focusing or localization errors can broaden vessels, duplicate vascular features, or blur adjacent structures. Incorporating the t-US-derived SoS map improved PA vessel converging, and reduced vessel duplication and blurring in PWD images. These improvements were also evident in ULM as sharper vascular features and clearer delineation of the microvasculature. These improvements are consistent with previous studies showing that acoustic heterogeneity can degrade both PA and US reconstruction and that incorporating more accurate SoS information can mitigate these effects [17, 34–36]. A practically important question is how much of this benefit requires a full SoS map rather than a simple SoS correction. We found that a dual-SoS model recovered partial improvement, with the t-US map adding further vessel separation and focusing in regions with stronger heterogeneity. Under the imaging conditions examined here, a substantial portion of the propagation-time error arose from the large SoS mismatch between the surrounding water and the animal body. Correctly identifying the body boundary and assigning separate SoS values to water and tissue therefore captured the global improvement, while spatial variations within the body provided regional corrections. From a practical perspective, dual-SoS reconstruction may offer a computationally simpler alternative when a reliable t-US-derived SoS map is not available.

The current SoS reconstruction method contains several approximations. We used first-arrival time-of-flight measurements and a straight-ray SART model, which assumes that acoustic waves propagate along undisturbed paths. In heterogeneous tissue, refraction causes the actual propagation paths to deviate from the straight lines connecting the Tx and Rx elements. Diffraction, multipath propagation, acoustic attenuation, and errors in first-arrival detection may introduce additional errors. Future implementations may incorporate refraction-corrected bent-ray tomography [49, 50] or full-wave inversion [51] to better account for heterogeneous acoustic propagation.

The r-US acquisition for B-mode, PWD, and ULM sequence was designed to balance image quality, frame rate, and data size. Coherent compounding of 20 plane-wave transmissions distributed around the full-ring aperture yielded a compounded frame rate of 526 Hz. A high frame rate of >500 Hz is important for ULM because excessive interframe MB displacement can disrupt trajectory linking, reduce the number of successfully reconstructed tracks, and bias velocity estimates, particularly in fast-flowing vessels [27, 39]. Increasing the number of transmitting angles can improve contrast-to-noise ratio, angular sampling, and the uniformity of the compounded point spread function [38]. With a fixed time interval between transmissions, however, the compounded frame rate decreases with the number of transmission angles, while the amount of RF data generated per frame increases. The selection of 20 transmission angles therefore represented a practical balance among image quality, frame rate, and data size.

The limited number of frames presented an additional challenge for ULM. To restrict the total imaging duration, only 3000 r-US frames were acquired at each scanning location. This number is relatively small for ULM, which requires the accumulation of sufficient localized MB events to sample the complete vascular network. Incomplete vascular sampling preferentially affects small, sparsely perfused, or out-of-plane vessels, and may bias vessel-density, diameter, and flow measurements toward large vessels. Respiratory-motion rejection further reduced the number of usable frames. Although the cross-correlation-based method removed motion-heavy frames, it did not compensate for moderate motion or local organ deformation. Discarding frames also introduced temporal discontinuities that could shorten MB trajectories and reduce tracking efficiency. Together, these factors limited the reliability of the reconstructed ULM microvascular networks. Future work may incorporate nonrigid image registration, motion-aware tracking, or respiratory gating to improve vascular sampling and trajectory reconstruction.

The photoswitching experiment demonstrated the benefit of combining molecular PA contrast with co-registered US vascular and anatomical information. Differential PA imaging isolated the DrBphP1-expressing tumor from endogenous background signals, while PWD and B-mode imaging localized the surrounding perfused vessels and tumor boundaries. Interleaved acquisition at 750 and 870 nm provided an estimate of tumor sO₂. However, these sO₂ measurements should be regarded as semiquantitative because wavelength-dependent optical fluence was not compensated. Multispectral acquisition using additional wavelengths, together with model- or measurement-based fluence correction, could improve the accuracy of sO₂ estimates. Repeated photoswitching cycles [41] and motion-aware fitting could also increase the sensitivity and specificity of the molecular detection.

In the renal IRI study, ULM-derived MB count and blood-volume measurements indicated an acute reduction in perfusion in the injured kidney at Day 1, followed by progressive recovery toward baseline by Day 13, whereas mean renal sO₂ remained reduced at later time points. This divergence suggests that restoration of detectable vascular perfusion may not necessarily indicate complete recovery of tissue oxygenation and may reflect persistent microvascular dysfunction, altered oxygen extraction, or metabolic changes after IRI. In contrast, mean vessel diameter and MB flow speed showed no clear longitudinal changes, although regional alterations may have been obscured by whole-kidney averaging, limited ULM sampling, and tracking bias toward larger or more consistently perfused vessels. The increased expression of *Ngal* and *Hif1a* in both kidneys and the stronger *Fabp1* elevation in the injured kidney further supported the presence of renal injury and systemic or compensatory responses after unilateral IRI. Nevertheless, these findings should be interpreted as proof of concept because the imaging cohort was small, sO₂ was estimated without fluence correction, and gene-expression measurements were obtained from a separate cohort. Larger longitudinal studies combining imaging with matched histology and renal-function biomarkers are needed to further understand these observations.

Overall, the integrated PA-US platform enables co-registered anatomical, vascular, hemodynamic, oxygenation, and molecular imaging within a unified framework. By combining t-US-based SoS correction, high-frame-rate r-US, ULM, and multispectral PACT, the system provides complementary measurements of tissue structures and functions and represents a versatile platform for multiparametric preclinical imaging.

## Acknowledgements

R.Y. thanks the support of the Fitzpatrick Foundation Scholars program in the Fitzpatrick Institute for Photonics (FIP) at Duke University. J.Y. thanks the support by the United States National Institutes of Health (NIH) grants RF1 NS115581, R01 NS111039, R01 EB028143, R01 DK139109, R01 DK052985, R01 MH135932, R01 ES036951; The United States National Science Foundation (NSF) CAREER award 2144788; Duke University Pratt Beyond the Horizon Grant; Eli Lilly Research Award Program; Chan Zuckerberg Initiative Grant (2020-226178 and 2024-349531); Duke University DST Spark Seed Grant; Duke Coulter Translational Grant; North Carolina Biotechnology Center Triangle Research Grant (2024-TRG-0041); American Heart Association Collaborative Science Award (25CSA1417550).

## Author contributions

J.Yao. conceived the study and led the study design, and supervised the research. R.Y. and T.V. designed and built the ring-array imaging platform. R.Y., H.H., N.W., T.V., and L.M. contributed to the image reconstruction and processing pipeline. R.Y. performed the acoustic field simulations. R.Y., J.Luo, X.C., and Y.X. performed the *in vivo* imaging experiments. J.Luo and X.C. contributed to tumor model establishment, animal preparations and fluorescence imaging. J.Li prepared the microbubbles. I.H. performed the IRI surgery and qPCR analyses. J.Luo and I.H. assisted with data analysis. I.H. and X.L provided pathological guidance. J.Yang, M.L., and P.S. provided ultrasound imaging resources. R.Y., I.H., J.Luo, and J.Yao. wrote the manuscript with input from all authors. Correspondence to Junjie Yao.

## Ethics declarations

### Competing interests

J.Yao, T.V., L.M. have a financial interest in Lumius Imaging, Inc., and J.Yao has a financial interest in Merge Labs, which did not support this work. The other authors declare no competing interests.

## Methods

### Data Acquisition and System Control

The ring-array PA-US imaging system (**Fig. 1a**) comprised two custom 256-element half-ring ultrasonic transducer arrays (Imasonics), each with a center frequency of 5 MHz, a transmit–receive −6-dB bandwidth of 3.5–6.5 MHz, and a receive-only −6-dB bandwidth of 0.15–7 MHz (**Supplementary Fig. 1**). Together, the two half-ring arrays formed a 512-element full-ring aperture with a diameter of 80 mm and an elevational radius of curvature of 37 mm. The array was multiplexed to a programmable 256- channel data acquisition system (Vantage 256, Verasonics), allowing up to 256 elements to be simultaneously activated for transmission or receiving. Received signals were digitized at 20.833 MS/s for PA and r-US acquisition and at 30 MS/s for t-US acquisition. For conventional PA imaging, optical excitation was provided by a Q-switched laser equipped with an optical parametric oscillator (OPO; SpitLight1200, Innolas) operating at a fixed pulse-repetition frequency (PRF) of 30 Hz. A power meter (Ophir) recorded the energy of each laser pulse for pulse-by-pulse energy normalization. A 635-nm CW diode laser was used only in photoswitching experiments.

For r-US acquisition, we employed a plane-wave transmission scheme (**Fig. 1b**). Each compounded r-US frame consisted of 20 plane-wave Tx/Rx events. For each event, a 64-element Tx aperture transmitted a single-cycle pulse at the center frequency with a peak-to-peak voltage of 10 V, while a 256-element Rx aperture located on the same side of the ring received the backscattered signals (**Supplementary Figs. 2a** and **2b**). The 20 Tx/Rx apertures were evenly distributed around the full 2π ring aperture, and consecutive events were separated by 95 μs. Thus, the acquisition time for one 20- angle compounded frame was 1.9 ms, corresponding to a frame rate of approximately 526 Hz.

Each r-US ensemble contained 250 consecutively acquired compounded frames, corresponding to 5,000 Tx/Rx events (**Fig. 1d**). The acquired data were initially stored in local memory on the Vantage acquisition modules and subsequently transferred by direct memory access (DMA) to a buffer in the host computer’s random-access memory (RAM) before acquisition of the next ensemble (**Fig. 1g**). Once the DMA transfer was completed and acquisition of the subsequent ensemble began, the preceding ensemble was transferred from the host RAM to disk. This data-handling strategy enabled sustained acquisition of large datasets while maintaining a fixed 95-μs interval between consecutive Tx/Rx events and a stable compounded frame rate of 526 Hz within each ensemble. The high, stable frame rate, together with the accumulation of consecutive frames, was necessary for reliable MB localization and trajectory reconstruction in ULM.

For t-US acquisition, each Tx event used a single array element to transmit a two-cycle pulse at the center frequency with a peak-to-peak voltage of 40 V (**Fig. 1c and Supplementary Fig. 2c**). The 256 elements on the opposite side of the ring served as the Rx aperture (**Supplementary Fig. 2d**). One t-US frame comprised 512 Tx/Rx events distributed around the entire ring aperture, with consecutive events separated by 200 μs (**Fig. 1f**). Each t-US ensemble contained three frames for improving signal to noise ratio, corresponding to a total of 1,536 Tx/Rx events. The data were transferred and saved after acquisition of each t-US ensemble was completed (**Fig. 1g**).

For PA acquisition, the 256-channel limitation required the 512 receive elements to be divided into two 256-element Rx apertures. Consequently, acquisition of one full-ring PA frame at a given wavelength required two consecutive laser pulses, one for each Rx aperture. This multiplexing reduced the effective full-ring PA frame rate from the 30-Hz laser PRF to 15 Hz. For multispectral acquisition, each wavelength was applied for two consecutive laser pulses before the OPO laser was switched to the next wavelength (**Fig. 1e**). The number of PA frames in each ensemble was selected according to the requirements of the individual experiment.

The lasers, data-acquisition system, and translation motor stage were triggered and synchronized using an FPGA-based control module (myRIO1900, National Instruments; **Fig. 1a**). Each scanning cycle consisted of the sequential acquisition and storage of one t-US ensemble, 12 r-US ensembles, and one PA ensemble, followed by translation of the stage to the next elevational position (**Fig. 1g**). Volumetric datasets were acquired over multiple scanning cycles, with the scanning step size selected according to the requirements of each experiment.

### t-US Image Reconstruction for SoS Estimation

The t-US data acquired during each scanning cycle were reconstructed into a 2D cross-sectional SoS map using first-arrival time-of-flight (TOF) estimation, followed by the simultaneous algebraic reconstruction technique (SART) under a straight-ray propagation model (**Fig. 2a**) [52, 53]. Although measurements were acquired over the full 2π angular aperture, only one measurement from each reciprocal Tx–Rx path pair was required for reconstruction because reciprocal propagation paths contain nominally redundant information. Reciprocal measurements were averaged before reconstruction to reduce measurement variability. For each Tx–Rx pair, the first-arrival time was identified within a predefined temporal window using an Akaike information criterion (AIC)-based first-motion picker [54]. The measured TOF was referenced to the expected propagation time through the homogeneous water background, whose SoS was determined from the water temperature. The resulting differential TOF represented the line integral of the slowness difference between the imaged object and the water background along each assumed straight propagation path. The spatial slowness distribution was reconstructed using SART with 40 iterations and subsequently converted to a SoS map. Reconstruction was performed within a circular ROI with a radius of 20 mm using a progressively refined spatial grid, yielding a final pixel size of 0.2 mm.

To reduce the influence of t-US reconstruction artifacts outside the animal, a binary body mask was applied to the reconstructed SoS map (**Supplementary Fig. 6**). The mask was generated from a US B-mode image reconstructed using the homogeneous water SoS, in which the boundary of the animal body could be readily identified and segmented. Reconstructed SoS values outside the body mask were replaced with the water SoS, whereas values within the mask were retained from the t-US reconstruction. The resulting masked SoS map was subsequently used to correct acoustic propagation times during PA and r-US reconstruction.

For the dual-SoS reconstructions evaluated in **Results**, the same body mask was used to divide the reconstruction domain into two homogeneous regions. The SoS was assigned a value of 1550 m/s within the animal body and 1521 m/s in the surrounding water at 36°C.

### r-US Image Reconstruction and Processing

r-US B-mode images were reconstructed using SoS-corrected DAS beamforming followed by coherent plane-wave compounding [38]. The RF data acquired from each transmission angle were first bandpass filtered according to the transducer bandwidth (**Fig. 2c**). Images were reconstructed on a grid with a pixel size of 70 μm. For each image pixel, the transmit and receive propagation times were calculated using the t-US-derived SoS map. The transmit propagation time was estimated by integrating the local slowness along the assumed straight path normal to the transmitted plane wavefront, whereas the receive propagation time was estimated along the straight path from the image pixel to each receiving element. The delayed RF signals were summed across the receive aperture, and the beamformed images from all 20 transmission angles were then coherently compounded. To accelerate reconstruction, DAS beamforming was implemented as GPU-accelerated sparse matrix multiplication [6]. A separate sparse reconstruction matrix was generated for each of the 20 plane-wave transmission angles. Each matrix encoded the SoS-corrected transmit and receive delays, RF-sample interpolation weights, and contributions from the active receiving elements. The matrices were reused for all r-US frames acquired at the same scanning position. When the translation stage advanced to a new scanning position, the matrices were recomputed using the corresponding updated t-US-derived SoS map. After reconstruction, each B-mode frame underwent a slight Gaussian smoothing (35) followed by 2D fast non-local means filtering [55] to reduce speckle-like intensity fluctuations.

### PWD and ULM Processing

To generate MB-enhanced PWD images, the reconstructed US B-mode frames strongly corrupted by respiratory motion were first identified within each 250-frame ensemble using cross-correlation and excluded from subsequent processing (**Fig. 2c**). On average, approximately 2⁄3 of the frames were retained after motion rejection.

Spatiotemporal singular value decomposition (SVD) clutter filtering was then applied to the motion-rejected ensemble to suppress tissue signals and isolate signals from circulating MBs [56]. The SVD cutoff was adaptively selected using the similarity of the spatial singular vectors [57]. Adaptive cutoff selection was used because MB concentration, tissue motion, and blood-flow characteristics varied across scanning positions, making a fixed cutoff suboptimal [58]. The PWD image was calculated by accumulating the signal power of the clutter-filtered images.

ULM processing was performed using the same clutter-filtered ensembles (**Fig. 2c**). Each clutter-filtered image was spatially interpolated to one-fifth of the original pixel spacing, corresponding to an interpolated spacing of 14 μm. MB centroids were localized by 2D normalized cross-correlation with a model 2D Gaussian point spread function. Candidate MB detections were retained when the normalized cross-correlation coefficient exceeded 0.5 and the normalized signal amplitude exceeded 0.05. To reduce spatial overlap among MB signals, signals associated with opposite flow directions were separated by Fourier-domain directional filtering and processed independently. The localized MB positions were linked across consecutive frames using the uTrack algorithm [59], which uses a linear-assignment framework for trajectory reconstruction. A maximum linking radius of 5 pixels was used, and a linear-motion Kalman filter was applied during trajectory reconstruction, with the maximum change in trajectory direction constrained to 45°. ULM intensity maps were generated by accumulating the localized MB positions, and blood flow velocity maps were calculated from the reconstructed MB trajectories.

### PA Image Reconstruction and Processing

The acquired PA signals were first deconvolved using the measured one-way receive impulse response of the transducer array (**Supplementary Fig. 1**) and subsequently bandpass filtered to suppress out-of-band noise (**Fig. 2b**). Images were reconstructed on a grid with a pixel size of 70 μm using SoS-corrected DAS beamforming. For each image pixel, the one-way acoustic propagation time to each receiving element was calculated by integrating the local slowness along the assumed straight propagation path through the t-US-derived SoS map. The appropriately delayed RF signals were then coherently summed across all 512 receiving elements to form a PA image. Again, because the array was multiplexed to a 256-channel data-acquisition system, each full-ring PA frame was assembled from two consecutive acquisition events, each containing signals from 256 receiving elements. Motion-corrupted PA frames were identified by cross-correlation and excluded before frame averaging to reduce respiratory-motion artifacts.

For blood oxygen saturation estimation, the RF signals acquired at 750 and 870 nm were normalized by the measured energy of the corresponding laser pulse before image reconstruction. The energy-normalized PA amplitudes at the two wavelengths were then linearly unmixed using the known molar extinction coefficients of oxyhemoglobin (HbO) and deoxyhemoglobin (Hb) [40, 60]. Blood oxygen saturation was subsequently calculated as HbO / HbO Hb . No wavelength-dependent optical fluence correction was applied.

### Animal Husbandry and Ethical Approval

BALB/c mice (8–10 weeks old) were used in all animal experiments. Founder mice were originally obtained from The Jackson Laboratory (Bar Harbor, ME, USA) and subsequently bred and maintained at the Duke University Division of Laboratory Animal Resources (DLAR).

Animals were housed in an AAALAC International-accredited specific pathogen-free (SPF) facility with ad libitum access to food and water under a 12 h light/12 h dark cycle. Both male and female mice were used as appropriate for the respective experiments. All animal procedures were approved by the Institutional Animal Care and Use Committee (IACUC) of Duke University (Protocols A203-22-12 and 60681) and were conducted in accordance with institutional guidelines and relevant regulations.

### Orthotopic Implantation of DrBphP1-4T1 Cells

The transgenic DrBphP1-4T1 cell line was obtained from an established cell line [61]. WT-4T1 cells and DrBphP1-4T1 cells were cultured in DMEM (Thermo Fisher Scientific) supplemented with 10% fetal bovine serum (FBS) and 1% penicillin–streptomycin at 37 °C in a humidified incubator with 5% CO₂ until reaching approximately 80–90% confluence. Before implantation, the cells were detached using trypsin, washed twice with phosphate-buffered saline (PBS), and resuspended in PBS. Immediately before injection, the cell suspension was mixed 1:1 (v/v) with growth factor-reduced Matrigel to obtain a final concentration of 2 × 10□cells/mL.

Female BALB/c mice (8–10 weeks old) were used for tumor implantation. Mice were anesthetized with 1.5% isoflurane in air delivered through a nose cone and placed in the supine position. The skin surrounding the right fourth mammary nipple was gently elevated using fine forceps to expose the mammary fat pad. A 27 G needle was inserted adjacent to the nipple into the fourth mammary fat pad, and 100 μL of the cell suspension (2 × 10□cells) was slowly injected. After injection, the needle was carefully withdrawn while the injection site was gently compressed with forceps for approximately 1 min to minimize reflux of the injected cell suspension. The contralateral fourth mammary fat pad was injected with an equal volume of WT-4T1 cells using the same procedure. Following imaging experiments, tumor-bearing mice were euthanized by carbon dioxide inhalation when the tumor diameter exceeded 1 cm.

### *In Vivo* IVIS Imaging

Fourteen days after orthotopic mammary fat pad implantation, mice were anesthetized with 1.5% isoflurane in air and imaged using an IVIS Kinetic imaging system. Fluorescence images were acquired in epi-fluorescence mode using a 665-nm excitation filter and a 750-nm emission filter. All animals were imaged using identical acquisition settings. Fluorescence images were processed using Living Image software and displayed as radiant efficiency in units of (photons s⁻¹ cm⁻² sr⁻¹)/(μW cm⁻²).

### PACT of the Photoswitchable Probe DrBphP1

To induce photoswitching of DrBphP1, the mouse was illuminated with 635-nm CW light for 6 s to switch the DrBphP1 molecules to the ON state (Fig. 4c). PA acquisition using interleaved 750- and 870-nm excitation began 5 s after the onset of CW illumination, resulting in a 1-s overlap between the 635-nm and pulsed OPO illumination. After the CW illumination was discontinued, PA acquisition continued for an additional 9.7 s, giving a total PA acquisition duration of approximately 10.7 s. Excitation at 750 nm switched DrBphP1 from the ON state to the OFF state while measuring its photoswitching response. The interleaved 870-nm excitation provided a second wavelength for estimating sO₂ together with the 750-nm measurements. PA signals were acquired only with the 750- and 870-nm excitation.

When photoswitching imaging was combined with US imaging, the multimodal acquisition sequence followed the scanning cycle shown in Fig. 1g. Each PA ensemble contained 320 laser-triggered acquisitions. Because each full-ring PA frame required two consecutive 256-element acquisitions, these acquisitions yielded 160 full-ring PA frames: 80 frames at 750 nm and 80 frames at 870 nm. The effective frame rate at each wavelength was 7.5 Hz.

For differential DrBphP1 imaging, the ON-state response was calculated by averaging the first 15 PA frames acquired at 750 nm, corresponding to the first 2.0 s of PA acquisition. The OFF-state response was calculated by averaging the remaining 65 frames acquired at 750 nm, corresponding to the subsequent 8.7 s. A differential image highlighting DrBphP1-specific contrast was generated by subtracting the mean OFF-state image from the mean ON-state image. For DrBphP1-specific imaging, the PA signals were low-pass filtered with a cutoff frequency of 1 MHz before image reconstruction to emphasize the low-frequency molecular signals. The motion-rejection procedure shown in Fig. 2b was not applied when generating the differential image.

### Kidney IRI Procedure

Mice (8–10 weeks old) were maintained under isoflurane for anesthesia and administered buprenorphine for pain control. They were then shaved and cleaned using betadine. With a right lateral incision, the right renal pedicle was exposed, ischemia was induced by placing a pedicle clamp for 30 minutes. Throughout the procedure, the mice were maintained at 36°C–37°C using a temperature-controlled heating pad. After 30 min, the clamp was removed to initiate reperfusion. The peritoneal and skin layers were closed separately using continuous transparent sutures selected to minimize PA signal and associated imaging artifacts.

### Data Analysis for Longitudinal IRI Imaging

Three-dimensional rectangular ROIs encompassing the right and left kidneys were manually defined for each imaging dataset. The following quantitative metrics were calculated separately within these ROIs. First, the right-to-left kidney MB count ratio was calculated by dividing the total number of MB localizations detected in the right-kidney ROI by that detected in the left-kidney ROI (Fig. 5d). The number of MB localizations was obtained by summing the values of all voxels in the corresponding ULM intensity map, in which each accumulated voxel value represented the number of localized MB events. Second, the blood volume ratio was calculated separately for each kidney as the volume occupied by detected perfused vessels divided by the total kidney ROI volume (Fig. 5e). The ULM intensity map was first binarized, and the blood volume ratio was calculated as the number of vessel-positive voxels divided by the total number of voxels within the kidney ROI. Third, vessel diameter was estimated from the binarized ULM intensity maps (Fig. 5f). The vascular centerlines were extracted using the MATLAB function bwskel with MinBranchLength set to 30 pixels. A Euclidean distance transform was then calculated from the binary vessel mask using bwdist. At each centerline pixel, the local vessel diameter was estimated as twice the distance to the nearest vessel boundary. The mean vessel diameter for each kidney was calculated by averaging the local diameter estimates within the corresponding ROI. Fourth, mean MB flow speed was calculated from the MB trajectories reconstructed during ULM processing (Fig. 5g). Instantaneous MB speed was determined from the interframe displacement of each linked MB localization divided by the frame interval. The mean whole-kidney MB flow speed was then calculated from the trajectory-derived speed measurements located within each kidney ROI. Finally, mean whole-kidney sO₂ was calculated from the PA-derived sO₂ values within each kidney ROI (Fig. 5h). Although the images in Fig. 5c display PA-derived sO₂ overlaid on PWD vascular images, PWD intensity was used only for visualization and was not used to calculate the whole-kidney mean sO₂.

### RNA Extraction and Semiquantitative Polymerase Chain Reaction (RT-qPCR)

RNA was extracted from tissue using Trizol Reagent (Invitrogen), followed by cDNA preparation using High-Capacity RNA-to-cDNA Kit (Applied Biosystems). Semiquantitative real-time PCR (ABI Prism 7500) was performed in triplicate using TaqMan master mix. We used the following TaqMan primers/probes: *Ngal*/*Lcn2*(Mm01324470_m1), *Hif1a* (Mm00468869_m1), *Fabp1* (Mm00444340_m1). We used the ddCt method to determine RNA expression, with *Gapdh* (Mm99999915_g1) serving as the internal control. Gene expression was normalized to the expression in naïve non-ischemic kidneys.

### Software

Image reconstruction and processing were performed using MATLAB R2025a (MathWorks). Acoustic field simulations were performed using the k-Wave toolbox v1.3 (www.k-wave.org) running on MATLAB. Data and statistical analyses were carried out in MATLAB and GraphPad Prism v10.

### Statistics and reproducibility

Data are presented as mean ± s.d. Sample numbers are provided in figure captions. Three independent samples were used for each experimental condition unless otherwise noted. Representative images were reproduced across independent experiments with similar results.

## Data availability

All data generated or analyzed during this study are included in the Article and its Supplementary Information.

## Notes

### Competing Interest Statement

The authors have declared no competing interest.

## References

[1] P. Beard, “Biomedical photoacoustic imaging,” Interface Focus, vol. 1, no. 4, pp. 602–631, 2011, doi: 10.1098/rsfs.2011.0028.

[2] L. V. Wang and J. Yao, “A practical guide to photoacoustic tomography in the life sciences,” Nature Methods, vol. 13, no. 8, pp. 627–638, 2016/08/01 2016, doi: 10.1038/nmeth.3925.

[3] N. Nyayapathi, E. Zheng, Q. Zhou, M. Doyley, and J. Xia, “Dual-modal photoacoustic and ultrasound imaging: from preclinical to clinical applications,” (in English), Frontiers in Photonics, Review vol. Volume 5 - 2024, 2024-February- 27 2024, doi: 10.3389/fphot.2024.1359784.

[4] Y. Yu, T. Feng, H. Qiu, Y. Gu, Q. Chen, C. Zuo, and H. Ma, “Simultaneous photoacoustic and ultrasound imaging: A review,” Ultrasonics, vol. 139, p. 107277, 2024/04/01/ 2024, doi: 10.1016/j.ultras.2024.107277.

[5] W. Choi, D. Oh, and C. Kim, “Practical photoacoustic tomography: Realistic limitations and technical solutions,” Journal of Applied Physics, vol. 127, no. 23, 2020, doi: 10.1063/5.0008401.

[6] V. Perrot, M. Polichetti, F. Varray, and D. Garcia, “So you think you can DAS? A viewpoint on delay-and-sum beamforming,” Ultrasonics, vol. 111, p. 106309, 2021/03/01/ 2021, doi: 10.1016/j.ultras.2020.106309.

[7] M. Xu and L. V. Wang, “Analytic explanation of spatial resolution related to bandwidth and detector aperture size in thermoacoustic or photoacoustic reconstruction,” Physical Review E, vol. 67, no. 5, p. 056605, 05/09/ 2003, doi: 10.1103/PhysRevE.67.056605.

[8] M. T. Rietberg, J. Gröhl, T. R. Else, S. E. Bohndiek, S. Manohar, and B. T. Cox, “Artifacts in photoacoustic imaging: Origins and mitigations,” Photoacoustics, vol. 45, p. 100745, 2025/10/01/ 2025, doi: 10.1016/j.pacs.2025.100745.

[9] S. Zhao, J. Hartanto, R. Joseph, C.-H. Wu, Y. Zhao, and Y.-S. Chen, “Hybrid photoacoustic and fast super-resolution ultrasound imaging,” Nature Communications, vol. 14, no. 1, p. 2191, 2023/04/18 2023, doi: 10.1038/s41467-023-37680-w.

[10] A. Garcia-Uribe, T. N. Erpelding, A. Krumholz, H. Ke, K. Maslov, C. Appleton, J. A. Margenthaler, and L. V. Wang, “Dual-Modality Photoacoustic and Ultrasound Imaging System for Noninvasive Sentinel Lymph Node Detection in Patients with Breast Cancer,” Scientific Reports, vol. 5, no. 1, p. 15748, 2015/10/29 2015, doi: 10.1038/srep15748.

[11] J. Robin, A. Özbek, M. Reiss, X. L. Dean-Ben, and D. Razansky, “Dual-Mode Volumetric Optoacoustic and Contrast Enhanced Ultrasound Imaging With Spherical Matrix Arrays,” IEEE Transactions on Medical Imaging, vol. 41, no. 4, pp. 846–856, 2022, doi: 10.1109/TMI.2021.3125398.

[12] Y. Tang, N. Wang, Z. Dong, M. Lowerison, A. d. Aguila, N. Johnston, T. Vu, C. Ma, Y. Xu, W. Yang, P. Song, and J. Yao, “Non-Invasive Deep-Brain Imaging With 3D Integrated Photoacoustic Tomography and Ultrasound Localization Microscopy (3D-PAULM),” IEEE Transactions on Medical Imaging, vol. 44, no. 2, pp. 994–1004, 2025, doi: 10.1109/TMI.2024.3477317.

[13] N. Wang, X. Yu, M. Lowerison, Q. Li, A. J. Canning, P. He, L. Dang, S. Degan, B. Mace, Y. Xu, R. Yao, J. Li, T. Zhou, J. Luo, B.-Z. Lin, D. A. Turner, X. Liu, D. Ta, J. Lovell, T. Vo-Dinh, W. Feng, P. Song, W. Yang, and J. Yao, “Noninvasive whole-brain imaging of glymphatic dynamics,” Science Advances, vol. 12, no. 25, p. eaee4926, 2026, doi: doi:10.1126/sciadv.aee4926.

[14] Y. Xu, R. Yao, H. Sheng, N. Wang, X. Yu, X. Cai, J. Cai, J. Luo, J. Li, W. Yang, P. Song, V. V. Verkhusha, and J. Yao, “3D-PAULM: Integrated Photoacoustic Tomography and Ultrasound Localization Microscopy for Multiparametric Brain and Tumor Imaging,” bioRxiv, p. 2026.04.30.722008, 2026, doi: 10.64898/2026.04.30.722008.

[15] E. Merčep, N. C. Burton, J. Claussen, and D. Razansky, “Whole-body live mouse imaging by hybrid reflection-mode ultrasound and optoacoustic tomography,” Optics Letters, vol. 40, no. 20, pp. 4643–4646, 2015/10/15 2015, doi: 10.1364/OL.40.004643.

[16] E. Merčep, J. L. Herraiz, X. L. Deán-Ben, and D. Razansky, “Transmission– reflection optoacoustic ultrasound (TROPUS) computed tomography of small animals,” Light: Science & Applications, vol. 8, no. 1, p. 18, 2019/01/30 2019, doi: 10.1038/s41377-019-0130-5.

[17] Y. Zhang and L. Wang, “Adaptive dual-speed ultrasound and photoacoustic computed tomography,” Photoacoustics, vol. 27, p. 100380, 2022/09/01/ 2022, doi: 10.1016/j.pacs.2022.100380.

[18] C. Lee, S. Cho, D. Lee, J. Lee, J.-I. Park, H.-J. Kim, S. H. Park, W. Choi, U. Kim, and C. Kim, “Panoramic volumetric clinical handheld photoacoustic and ultrasound imaging,” Photoacoustics, vol. 31, p. 100512, 2023/06/01/ 2023, doi: 10.1016/j.pacs.2023.100512.

[19] L. Menozzi, T. Vu, A. J. Canning, H. Rawtani, C. Taboada, M. E. Abi Antoun, C. Ma, J. Delia, V. T. Nguyen, S.-W. Cho, J. Chen, T. Charity, Y. Xu, P. Tran, J. Xia, G. M. Palmer, T. Vo-Dinh, L. Feng, and J. Yao, “Three-dimensional diffractive acoustic tomography,” Nature Communications, vol. 16, no. 1, p. 1149, 2025/01/29 2025, doi: 10.1038/s41467-025-56435-3.

[20] S. Zhao, X. Zhang, K. Bailey, S. Pai, Y. Zhao, and Y.-S. Chen, “Label-Free Dual-Modal Photoacoustic/Ultrasound Localization Imaging for Studying Acute Kidney Injury,” Advanced Science, vol. 12, no. 22, p. 2414306, 2025, doi: 10.1002/advs.202414306.

[21] L. Zhang, F. Meng, X. Zu, L. Luo, W. Ling, Y. Cai, S. Dai, B. Zhai, G. Lyu, and Q. Wu, “Dual-modal photoacoustic and ultrasound imaging for early diagnosis of ovarian torsion and evaluation of long-term tissue hypoxia,” Scientific Reports, vol. 15, no. 1, p. 40675, 2025/11/19 2025, doi: 10.1038/s41598-025-17381-8.

[22] Y. Zhang and L. Wang, “Video-Rate Ring-Array Ultrasound and Photoacoustic Tomography,” IEEE Transactions on Medical Imaging, vol. 39, no. 12, pp. 4369–4375, 2020, doi: 10.1109/TMI.2020.3017815.

[23] J. Pan, Q. Li, Y. Feng, R. Zhong, Z. Fu, S. Yang, W. Sun, B. Zhang, Q. Sui, J. Chen, Y. Shen, and Z. Li, “Parallel interrogation of the chalcogenide-based micro-ring sensor array for photoacoustic tomography,” Nature Communications, vol. 14, no. 1, p. 3250, 2023/06/05 2023, doi: 10.1038/s41467-023-39075-3.

[24] G. Wissmeyer, M. A. Pleitez, A. Rosenthal, and V. Ntziachristos, “Looking at sound: optoacoustics with all-optical ultrasound detection,” Light: Science & Applications, vol. 7, 12/01 2018, doi: 10.1038/s41377-018-0036-7.

[25] N. T. Huynh, E. Zhang, O. Francies, F. Kuklis, T. Allen, J. Zhu, O. Abeyakoon, F. Lucka, M. Betcke, J. Jaros, S. Arridge, B. Cox, A. A. Plumb, and P. Beard, “A fast all-optical 3D photoacoustic scanner for clinical vascular imaging,” Nature Biomedical Engineering, vol. 9, no. 5, pp. 638–655, 2025/05/01 2025, doi: 10.1038/s41551-024-01247-x.

[26] O. Ogunlade, R. Ellwood, E. Zhang, B. T. Cox, and P. Beard, “Three-Dimensional Whole-Body Small Animal Photoacoustic Tomography Using a Multi-View Fabry-Perot Scanner,” IEEE Transactions on Medical Imaging, vol. 44, no. 4, pp. 1922–1930, 2025, doi: 10.1109/TMI.2024.3522220.

[27] C. Errico, J. Pierre, S. Pezet, Y. Desailly, Z. Lenkei, O. Couture, and M. Tanter, “Ultrafast ultrasound localization microscopy for deep super-resolution vascular imaging,” Nature, vol. 527, no. 7579, pp. 499–502, 2015/11/01 2015, doi: 10.1038/nature16066.

[28] J. Yang, S. Choi, and C. Kim, “Practical review on photoacoustic computed tomography using curved ultrasound array transducer,” Biomedical Engineering Letters, vol. 12, no. 1, pp. 19–35, 2022/02/01 2022, doi: 10.1007/s13534-021-00214-8.

[29] X. L. Deán-Ben and D. Razansky, “On the link between the speckle free nature of optoacoustics and visibility of structures in limited-view tomography,” Photoacoustics, vol. 4, no. 4, pp. 133–140, 2016/12/01/ 2016, doi: 10.1016/j.pacs.2016.10.001.

[30] Q. Huang and Z. Zeng, “A Review on Real-Time 3D Ultrasound Imaging Technology,” BioMed Research International, vol. 2017, no. 1, p. 6027029, 2017, doi: 10.1155/2017/6027029.

[31] D. H. Turnbull and F. S. Foster, “Fabrication and characterization of transducer elements in two-dimensional arrays for medical ultrasound imaging,” IEEE Transactions on Ultrasonics, Ferroelectrics, and Frequency Control, vol. 39, no. 4, pp. 464–475, 1992, doi: 10.1109/58.148536.

[32] J. Xia, M. Chatni, K. Maslov, Z. Guo, K. Wang, M. Anastasio, and L. Wang, “Whole-body ring-shaped confocal photoacoustic computed tomography of small animals in vivo,” Journal of Biomedical Optics, vol. 17, no. 5, p. 050506, 2012. [Online]. Available: 10.1117/1.JBO.17.5.050506.

[33] G. Li, J. Xia, K. Wang, K. Maslov, M. A. Anastasio, and L. V. Wang, “Tripling the detection view of high-frequency linear-array-based photoacoustic computed tomography by using two planar acoustic reflectors,” Quantitative Imaging in Medicine and Surgery, vol. 5, no. 1, pp. 57–62, 2014. [Online]. Available: https://qims.amegroups.org/article/view/5018.

[34] L. Li, L. Zhu, C. Ma, L. Lin, J. Yao, L. Wang, K. Maslov, R. Zhang, W. Chen, J. Shi, and L. V. Wang, “Single-impulse panoramic photoacoustic computed tomography of small-animal whole-body dynamics at high spatiotemporal resolution,” Nature Biomedical Engineering, vol. 1, no. 5, p. 0071, 2017/05/10 2017, doi: 10.1038/s41551-017-0071.

[35] M. Cui, H. Zuo, X. Wang, K. Deng, J. Luo, and C. Ma, “Adaptive photoacoustic computed tomography,” Photoacoustics, vol. 21, p. 100223, 2021/03/01/ 2021, doi: 10.1016/j.pacs.2020.100223.

[36] J. Xia, C. Huang, K. Maslov, M. A. Anastasio, and L. V. Wang, “Enhancement of photoacoustic tomography by ultrasonic computed tomography based on optical excitation of elements of a full-ring transducer array,” Optics Letters, vol. 38, no. 16, pp. 3140–3143, 2013/08/15 2013, doi: 10.1364/OL.38.003140.

[37] T. Vu, P. Klippel, A. J. Canning, C. Ma, H. Zhang, L. A. Kasatkina, Y. Tang, J. Xia, V. V. Verkhusha, T. Vo-Dinh, Y. Jing, and J. Yao, “On the Importance of Low-Frequency Signals in Functional and Molecular Photoacoustic Computed Tomography,” IEEE Transactions on Medical Imaging, vol. 43, no. 2, pp. 771–783, 2024, doi: 10.1109/TMI.2023.3320668.

[38] G. Montaldo, M. Tanter, J. Bercoff, N. Benech, and M. Fink, “Coherent plane-wave compounding for very high frame rate ultrasonography and transient elastography,” *IEEE Transactions on Ultrasonics*, Ferroelectrics, and Frequency Control, vol. 56, no. 3, pp. 489–506, 2009, doi: 10.1109/TUFFC.2009.1067.

[39] X. Guo, D. Ta, and K. Xu, “Frame rate effects and their compensation on super-resolution microvessel imaging using ultrasound localization microscopy,” Ultrasonics, vol. 132, p. 107009, 2023/07/01/ 2023, doi: 10.1016/j.ultras.2023.107009.

[40] S. L. Jacques, “Optical properties of biological tissues: a review,” Physics in Medicine & Biology, vol. 58, no. 11, p. R37, 2013/05/10 2013, doi: 10.1088/0031-9155/58/11/R37.

[41] J. Yao, A. A. Kaberniuk, L. Li, D. M. Shcherbakova, R. Zhang, L. Wang, G. Li, V. V. Verkhusha, and L. V. Wang, “Multiscale photoacoustic tomography using reversibly switchable bacterial phytochrome as a near-infrared photochromic probe,” Nature Methods, vol. 13, no. 1, pp. 67–73, 2016/01/01 2016, doi: 10.1038/nmeth.3656.

[42] E. E. Hesketh, A. Czopek, M. Clay, G. Borthwick, D. Ferenbach, D. Kluth, and J. Hughes, Renal Ischaemia Reperfusion Injury: A Mouse Model of Injury and Regeneration (no. 88). 1940-087X, 2014, p. e51816.

[43] S. Frydman, O. Freund, L. Zornitzki, H. A. Katash, S. Banai, and Y. Shacham, “Indexed neutrophil gelatinase associated lipocalin: a novel biomarker for the assessment of acute kidney injury,” Journal of Nephrology, vol. 37, no. 2, pp. 401–407, 2024/03/01 2024, doi: 10.1007/s40620-023-01800-y.

[44] H. Thiessen-Philbrook, J. Koyner, M. Shlipak, R. Kim, S. Coca, C. Edelstein, C. Parikh, M. Zappitelli, K. Sint, A. Garg, C. Krawczeski, S. Li, and P. Devarajan, “Postoperative Biomarkers Predict Acute Kidney Injury and Poor Outcomes after Pediatric Cardiac Surgery,” Journal of the American Society of Nephrology, vol. 22, no. 9, pp. 1737–1747, 2011/09/01 2011, doi: 10.1681/ASN.2010111163.

[45] P. Kang and F. Cheng, “Mechanisms and therapeutic prospects of hypoxia-inducible factor 1-alpha in acute kidney injury: a systematic review,” (in English), Frontiers in Cell and Developmental Biology, Review vol. Volume 13 - 2025, 2026-January-12 2026, doi: 10.3389/fcell.2025.1660433.

[46] M. Matsuo, K. Taguchi, Y. Yokota, K. Fukami, and T. Igawa, “The impact of ischemic reperfusion injury on contralateral kidneys and the determinants of renal prognosis after robot-assisted partial nephrectomy,” (in eng), PLoS One, vol. 20, no. 4, p. e0321769, 2025, doi: 10.1371/journal.pone.0321769.

[47] A. J. Polichnowski, K. A. Griffin, H. Licea-Vargas, R. Lan, M. M. Picken, J. Long, G. A. Williamson, C. Rosenberger, S. Mathia, M. A. Venkatachalam, and A. K. Bidani, “Pathophysiology of unilateral ischemia-reperfusion injury: importance of renal counterbalance and implications for the AKI-CKD transition,” American Journal of Physiology-Renal Physiology, vol. 318, no. 5, pp. F1086–F1099, 2020, doi: 10.1152/ajprenal.00590.2019.

[48] H. Li, E. E. Dixon, H. Wu, and B. D. Humphreys, “Comprehensive single-cell transcriptional profiling defines shared and unique epithelial injury responses during kidney fibrosis,” Cell Metabolism, vol. 34, no. 12, pp. 1977–1998.e9, 2022, doi: 10.1016/j.cmet.2022.09.026.

[49] S. Li, M. Jackowski, D. P. Dione, T. Varslot, L. H. Staib, and K. Mueller, “Refraction corrected transmission ultrasound computed tomography for application in breast imaging,” (in eng), Med Phys, vol. 37, no. 5, pp. 2233–46, May 2010, doi: 10.1118/1.3360180.

[50] A. Javaherian, F. Lucka, and B. T. Cox, “Refraction-corrected ray-based inversion for three-dimensional ultrasound tomography of the breast,” Inverse Problems, vol. 36, no. 12, p. 125010, 2020/12/14 2020, doi: 10.1088/1361-6420/abc0fc.

[51] R. Ali, T. M. Mitcham, T. Brevett, Ò. C. Agudo, C. D. Martinez, C. Li, M. M. Doyley, and N. Duric, “2-D Slicewise Waveform Inversion of Sound Speed and Acoustic Attenuation for Ring Array Ultrasound Tomography Based on a Block LU Solver,” IEEE Transactions on Medical Imaging, vol. 43, no. 8, pp. 2988–3000, 2024, doi: 10.1109/TMI.2024.3383816.

[52] A. H. Andersen and A. C. Kak, “Simultaneous Algebraic Reconstruction Technique (SART): A superior implementation of the ART algorithm,” Ultrasonic Imaging, vol. 6, no. 1, pp. 81–94, 1984/01/01/ 1984, doi: 10.1016/0161-7346(84)90008-7.

[53] A. C. Kak and A. H. Andersen, Principles of Computerized Tomographic Imaging. IEEE Press, 1988.

[54] C. Li, L. Huang, N. Duric, H. Zhang, and C. Rowe, “An improved automatic time-of-flight picker for medical ultrasound tomography,” Ultrasonics, vol. 49, no. 1, pp. 61–72, 2009/01/01/ 2009, doi: 10.1016/j.ultras.2008.05.005.

[55] J. Darbon, A. Cunha, T. F. Chan, S. Osher, and G. J. Jensen, “Fast nonlocal filtering applied to electron cryomicroscopy,” in *2008 5th IEEE International Symposium on Biomedical Imaging: From Nano to Macro*, 14-17 May 2008 2008, pp. 1331–1334, doi: 10.1109/ISBI.2008.4541250.

[56] C. Demené, T. Deffieux, M. Pernot, B. F. Osmanski, V. Biran, J. L. Gennisson, L. A. Sieu, A. Bergel, S. Franqui, J. M. Correas, I. Cohen, O. Baud, and M. Tanter, “Spatiotemporal Clutter Filtering of Ultrafast Ultrasound Data Highly Increases Doppler and fUltrasound Sensitivity,” IEEE Transactions on Medical Imaging, vol. 34, no. 11, pp. 2271–2285, 2015, doi: 10.1109/TMI.2015.2428634.

[57] J. Baranger, B. Arnal, F. Perren, O. Baud, M. Tanter, and C. Demené, “Adaptive Spatiotemporal SVD Clutter Filtering for Ultrafast Doppler Imaging Using Similarity of Spatial Singular Vectors,” IEEE Transactions on Medical Imaging, vol. 37, no. 7, pp. 1574–1586, 2018, doi: 10.1109/TMI.2018.2789499.

[58] P. Song, A. Manduca, J. D. Trzasko, and S. Chen, “Ultrasound Small Vessel Imaging With Block-Wise Adaptive Local Clutter Filtering,” IEEE Transactions on Medical Imaging, vol. 36, no. 1, pp. 251–262, 2017, doi: 10.1109/TMI.2016.2605819.

[59] P. Roudot, W. R. Legant, Q. Zou, K. M. Dean, T. Isogai, E. S. Welf, A. F. David, D. W. Gerlich, R. Fiolka, E. Betzig, and G. Danuser, “u-track3D: Measuring, navigating, and validating dense particle trajectories in three dimensions,” Cell Reports Methods, vol. 3, no. 12, p. 100655, 2023/12/18/ 2023, doi: 10.1016/j.crmeth.2023.100655.

[60] M. Li, Y. Tang, and J. Yao, “Photoacoustic tomography of blood oxygenation: A mini review,” Photoacoustics, vol. 10, pp. 65–73, 2018/06/01/ 2018, doi: 10.1016/j.pacs.2018.05.001.

[61] L. A. Kasatkina, C. Ma, H. Sheng, M. Lowerison, L. Menozzi, M. Baloban, Y. Tang, Y. Xu, L. Humayun, T. Vu, P. Song, J. Yao, and V. V. Verkhusha, “Deep-tissue high-sensitivity multimodal imaging and optogenetic manipulation enabled by biliverdin reductase knockout,” Nature Communications, vol. 16, no. 1, p. 6469, 2025/07/14 2025, doi: 10.1038/s41467-025-61532-4.

