## supplementary document for "Whole-body Super-resolution Functional and Molecular Imaging with Panoramic Photoacoustic–Ultrasound Tomography"

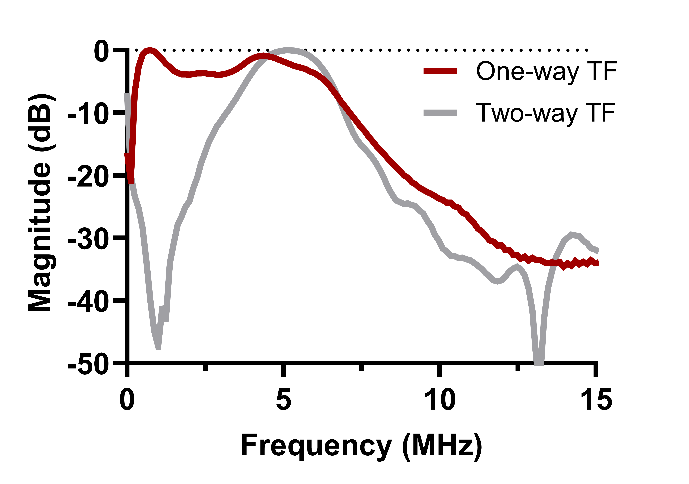

**Supplementary Figure 1*.* Frequency responses of the ring-array transducer*.*** Normalized one-way receiving and two-way transmit–receiving transfer function (TF) magnitudes of the ring-array transducer.

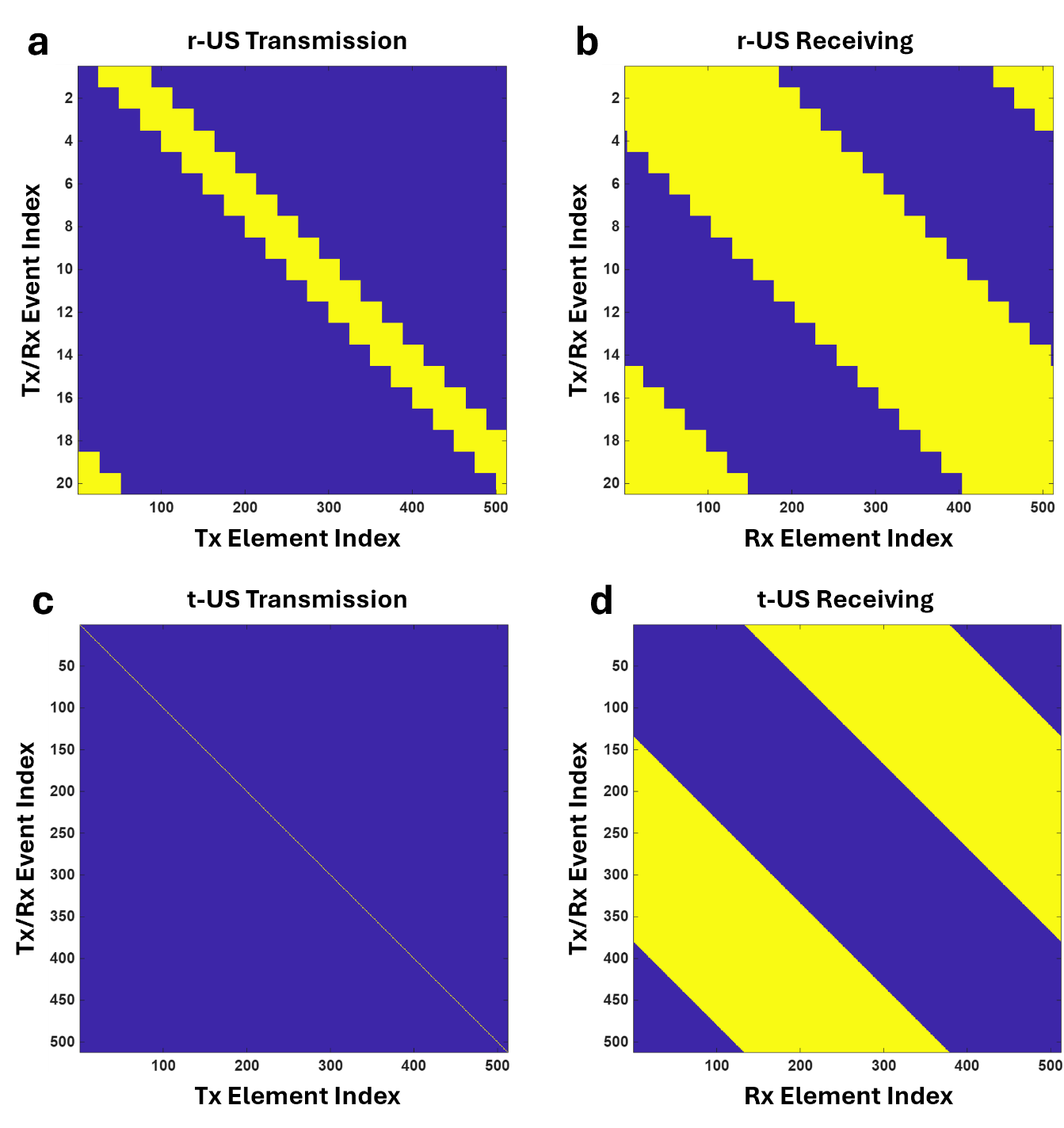

**Supplementary Figure 2. Transmission (Tx) and receiving (Rx) aperture configurations for reflection-mode ultrasound (r-US) and transmission-mode ultrasound (t-US)** **acquisition.** Active array elements in each Tx/Rx event are shown in yellow as a function of element index and event number. **(a, b)** r-US configuration showing **(a)** the 64-element Tx apertures and **(b)** the corresponding 256-element Rx apertures. The Tx and Rx apertures were positioned on the same side of the ring array and shared the same center point. Twenty plane-wave Tx/Rx events were evenly distributed around the full $2\pi$ aperture. **(c, d)** t-US configuration showing **(c)** the single-element Tx apertures and **(d)** the corresponding 256-element Rx apertures positioned on the opposite side of the ring array. A complete t-US frame contained 512 Tx/Rx events, with each array element sequentially serving as the transmitter.

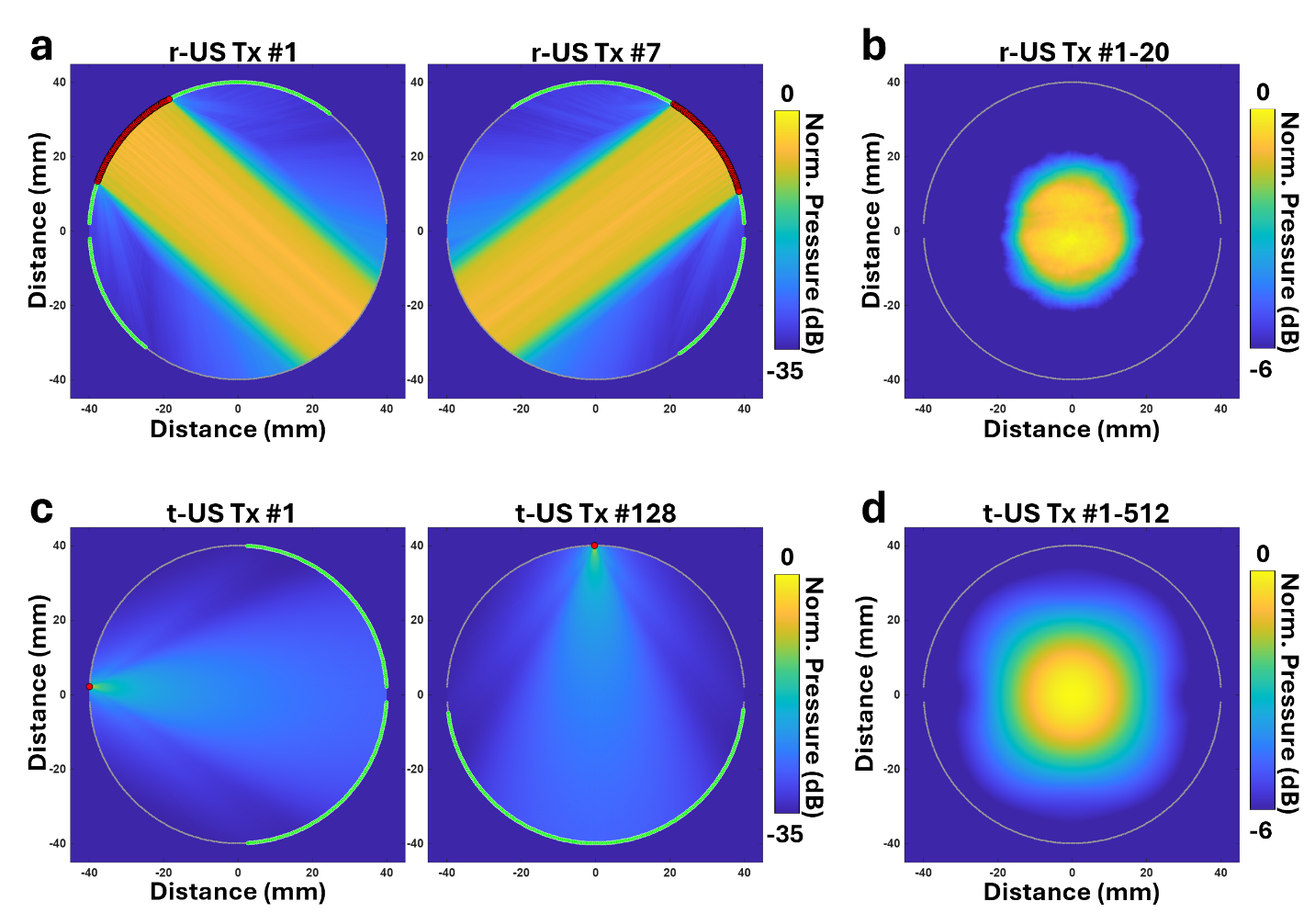

**Supplementary Figure 3. Simulated 2D acoustic pressure fields for r-US and t-US transmission.** Active Tx elements are shown in red, and the corresponding Rx elements are shown in green. Pressure magnitudes are normalized and displayed in decibels. **(a)** Representative pressure fields generated by r-US plane-wave Tx event 1 and 7. **(b)** Cumulative pressure field produced by all 20 r-US Tx events. **(c)** Representative pressure fields generated by t-US single-element Tx event 1 and 128. **(d)** Cumulative pressure field produced by all 512 t-US Tx events. Simulations were performed using the k-Wave toolbox in MATLAB with a spatial grid spacing of 0.1 mm. The simulated array geometry, including element positions, element widths, and Tx/Rx aperture configurations, was identical to the experimental system.

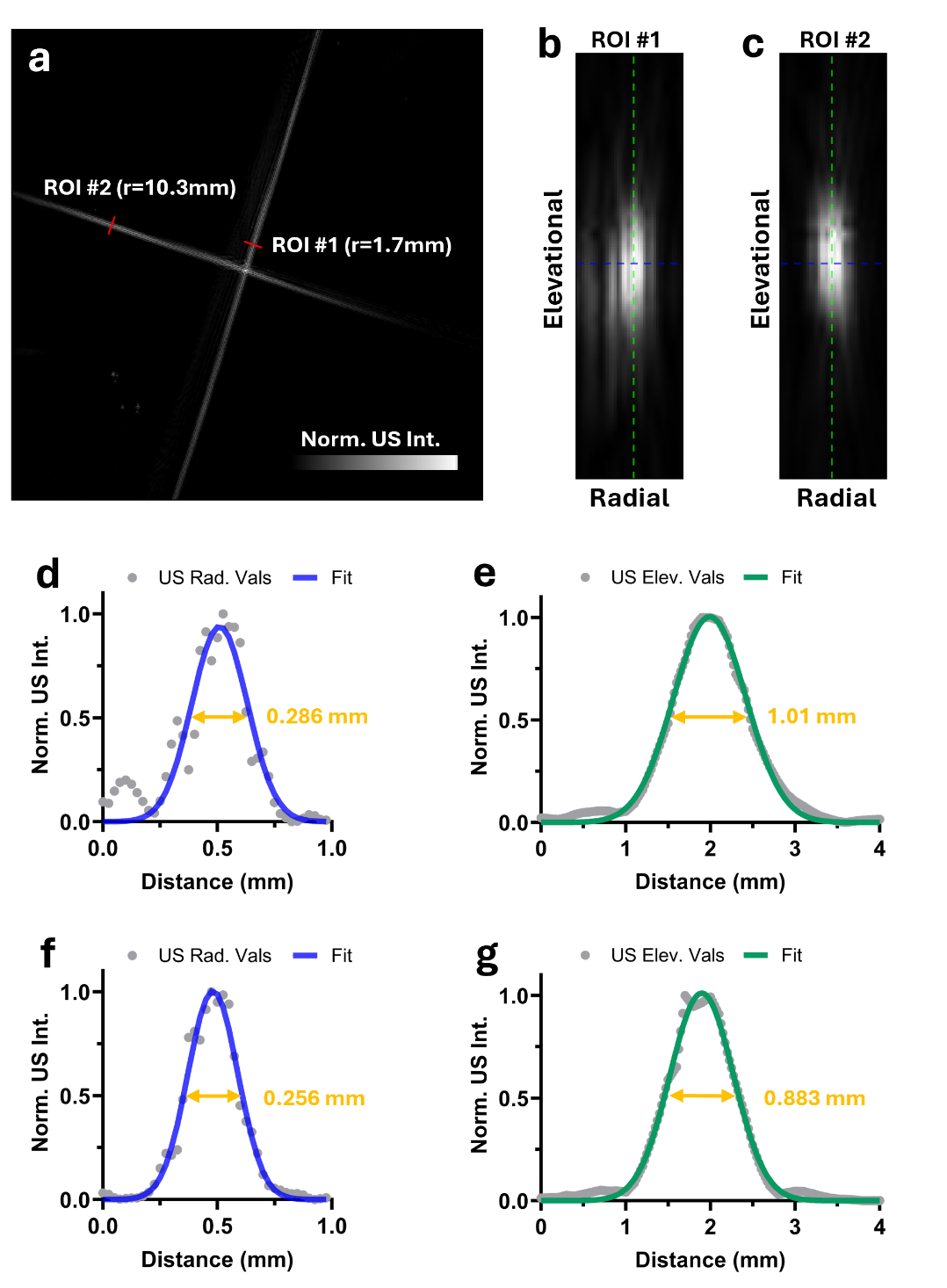

**Supplementary Figure 4. Spatial-resolution characterization of US B-mode imaging using a wire phantom.** **(a)** B-mode image of two orthogonal copper wires with a nominal diameter of 10 μm. Two regions of interest were selected at radial distances of 1.7 mm (ROI #1) and 10.3 mm (ROI #2) from the center of the ring-array transducer. **(b, c)** Radial–elevational cross-sectional images of **(b)** ROI #1 and **(c)** ROI #2. The blue and green dashed lines indicate the radial and elevational profile locations, respectively. **(d, e)** Normalized radial and elevational profiles from ROI #1, respectively. FWHM values of 0.286 mm in the radial direction and 1.01 mm in the elevational direction were obtained from the fitted profiles. **(f, g)** Normalized radial and elevational profiles from ROI #2, respectively, yielding FWHM values of 0.256 mm in the radial direction and 0.883 mm in the elevational direction. The US resolutions are similar across the field of view of the ring array.

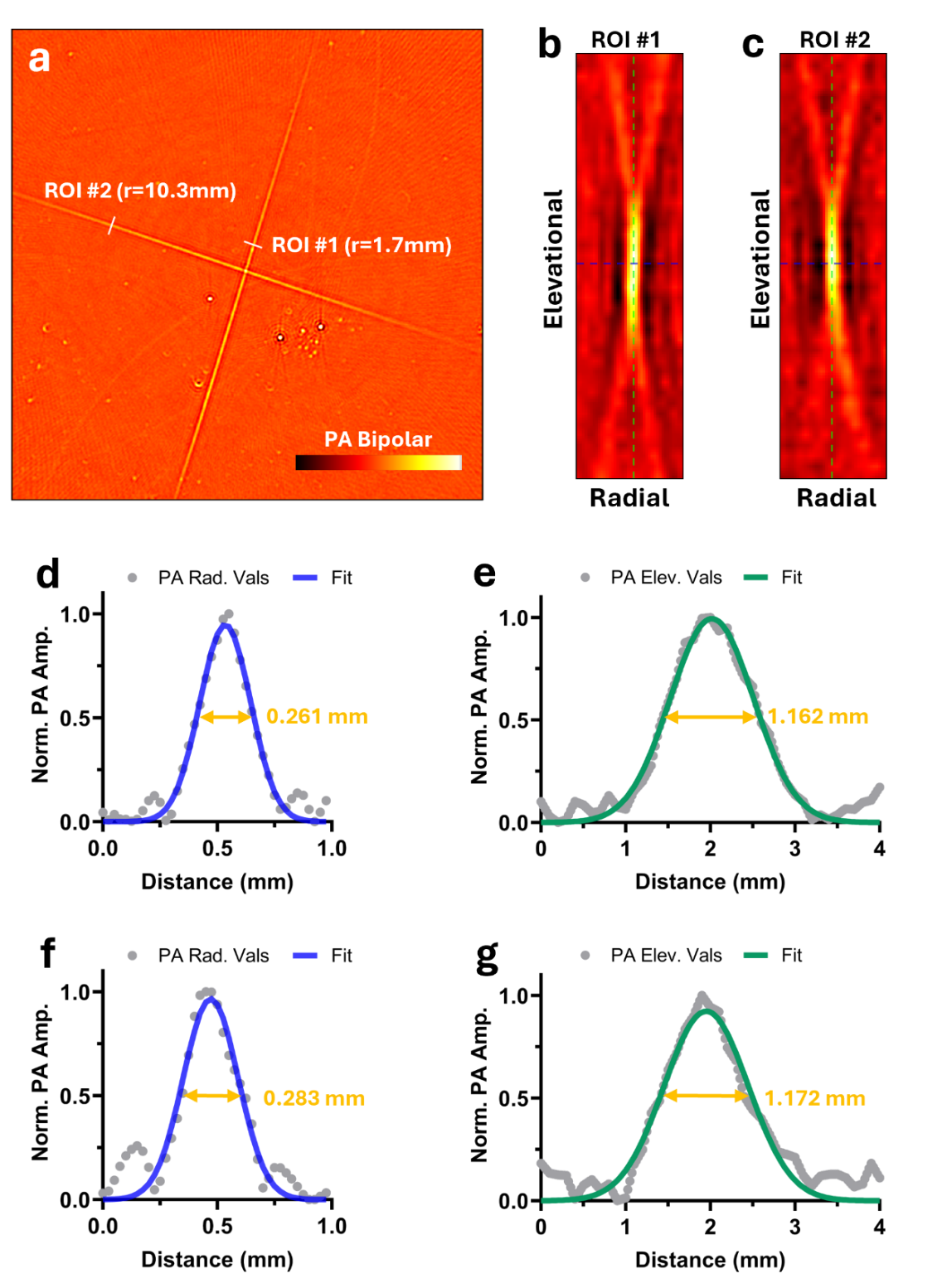

**Supplementary Figure 5. Spatial-resolution characterization of PA imaging using a wire phantom.** **(a)** Bipolar PA image of two orthogonal copper wires with a nominal diameter of 10 μm, acquired at 1064-nm excitation. Two regions of interest were selected at radial distances of 1.7 mm (ROI #1) and 10.3 mm (ROI #2) from the center of the ring-array transducer. **(b, c)** Radial–elevational cross-sectional PA images of **(b)** ROI #1 and **(c)** ROI #2. The blue and green dashed lines indicate the radial and elevational profile locations, respectively. **(d, e)** Normalized radial and elevational envelope profiles from ROI #1, respectively. FWHM values of 0.261 mm in the radial direction and 1.162 mm in the elevational direction were obtained from the fitted profiles. **(f, g)** Normalized radial and elevational envelope profiles from ROI #2, respectively, yielding FWHM values of 0.283 mm in the radial direction and 1.172 mm in the elevational direction. The PA resolutions are similar across the field of view of the ring array. Although the images in panels **(a–c)** are displayed using bipolar PA amplitudes, the line profiles were converted to analytic-signal envelopes using a Hilbert transform before profile fitting and FWHM measurement for a fair comparison with US B-mode.

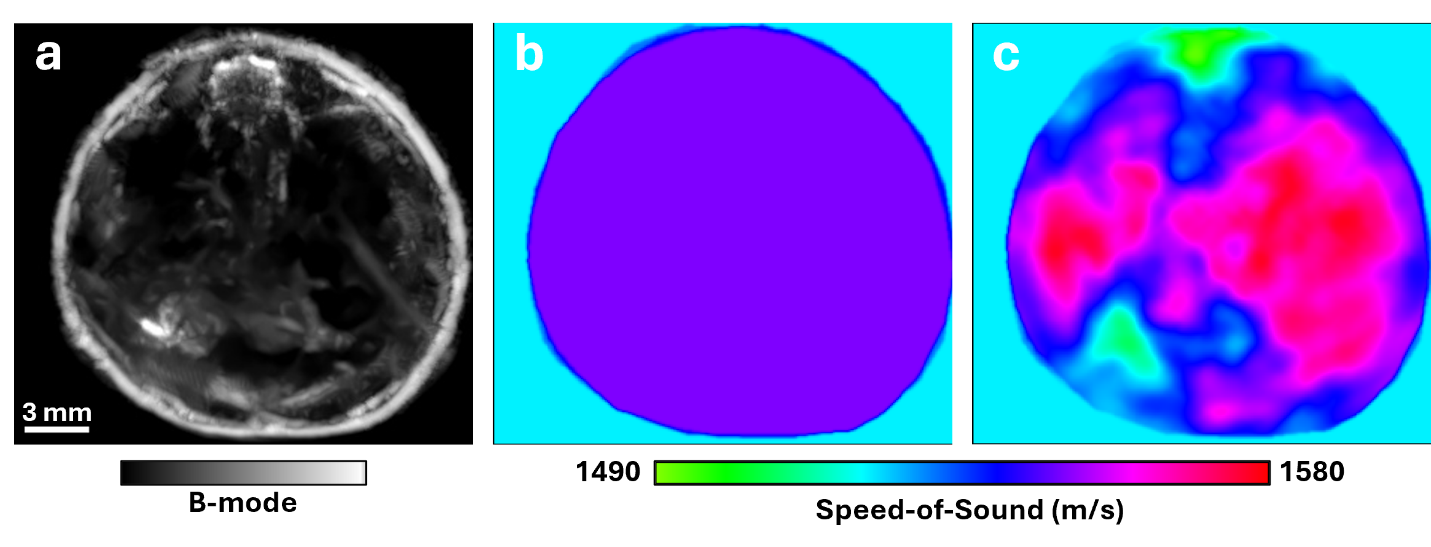

**Supplementary Figure 6. Dual-SoS map and spatially varying SoS map.** **(a)** B-mode cross-section image at the same location as the PA and PWD images shown in **Figs. 2d–2g**. A binary body mask was generated by segmenting the skin surface of the animal in the B-mode image. **(b)** Dual-SoS map generated by assigning a uniform SoS of 1550 m/s within the body and 1521 m/s outside the body. **(c)** Masked t-US-derived SoS map. The t-US SoS values were retained within the body, whereas values outside the body were assigned with 1521 m/s, corresponding to the SoS of the 36 °C water bath.

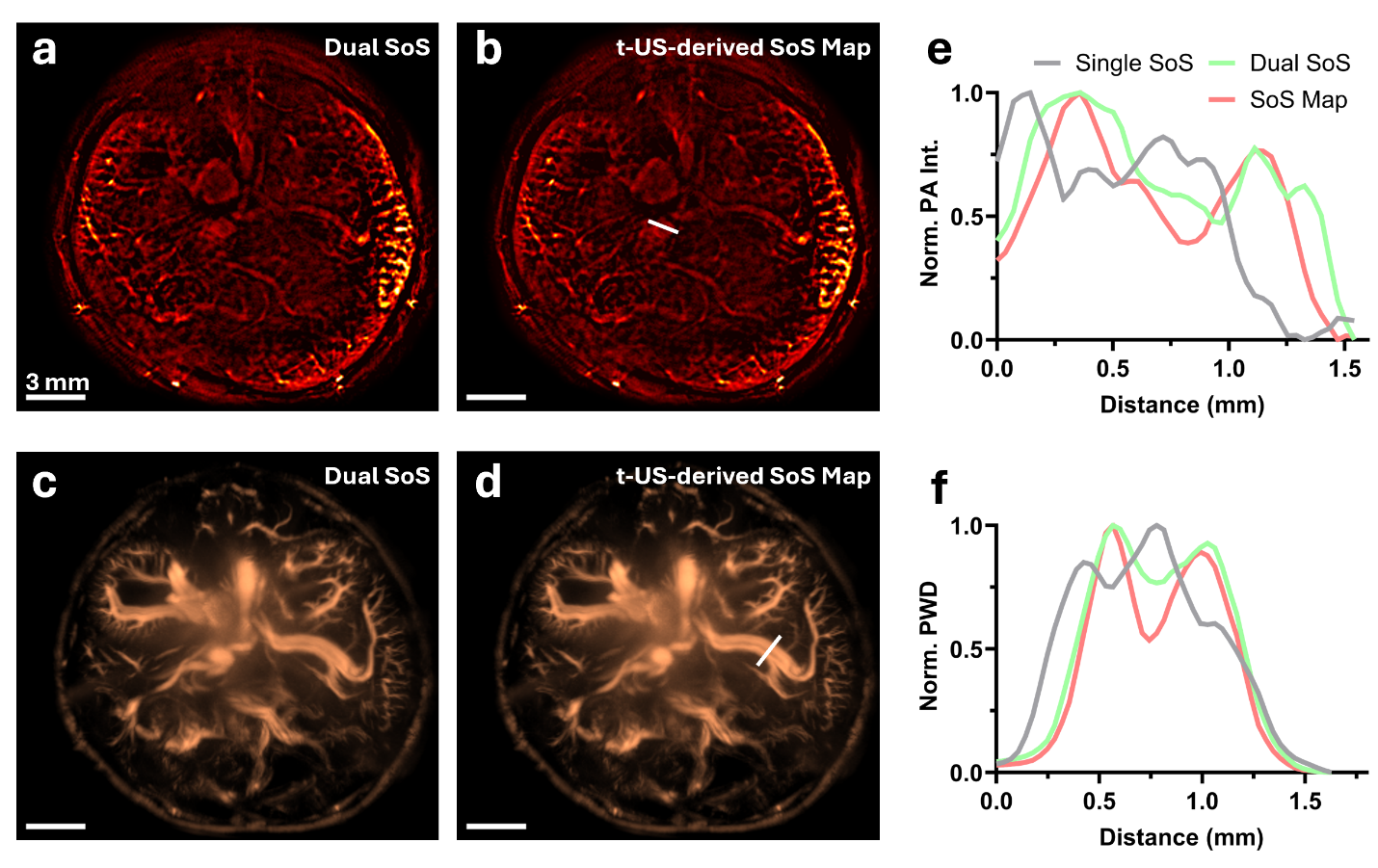

**Supplementary Figure 7. Comparison of single-SoS-, dual-SoS-, and spatially-varying SoS-map-assisted reconstruction.** **(a, b)** PA images acquired at 1064 nm and reconstructed using **(a)** a dual-SoS map or **(b)** a spatially varying t-US-derived SoS map. **(c, d)** Corresponding r-US PWD images reconstructed using **(c)** the dual-SoS map or **(d)** the t-US-derived SoS map. **(e, f)** Normalized line profiles extracted from the locations indicated by the white lines in **(b)** and **(d),** respectively. Profiles from reconstruction using a single SoS of 1521 m/s are included for comparison. Both dual-SoS and spatially-varying SoS-map-assisted reconstruction improved vessel converging relative to single-SoS reconstruction, while the spatially-varying SoS map provided better vessel separation in the selected regions.

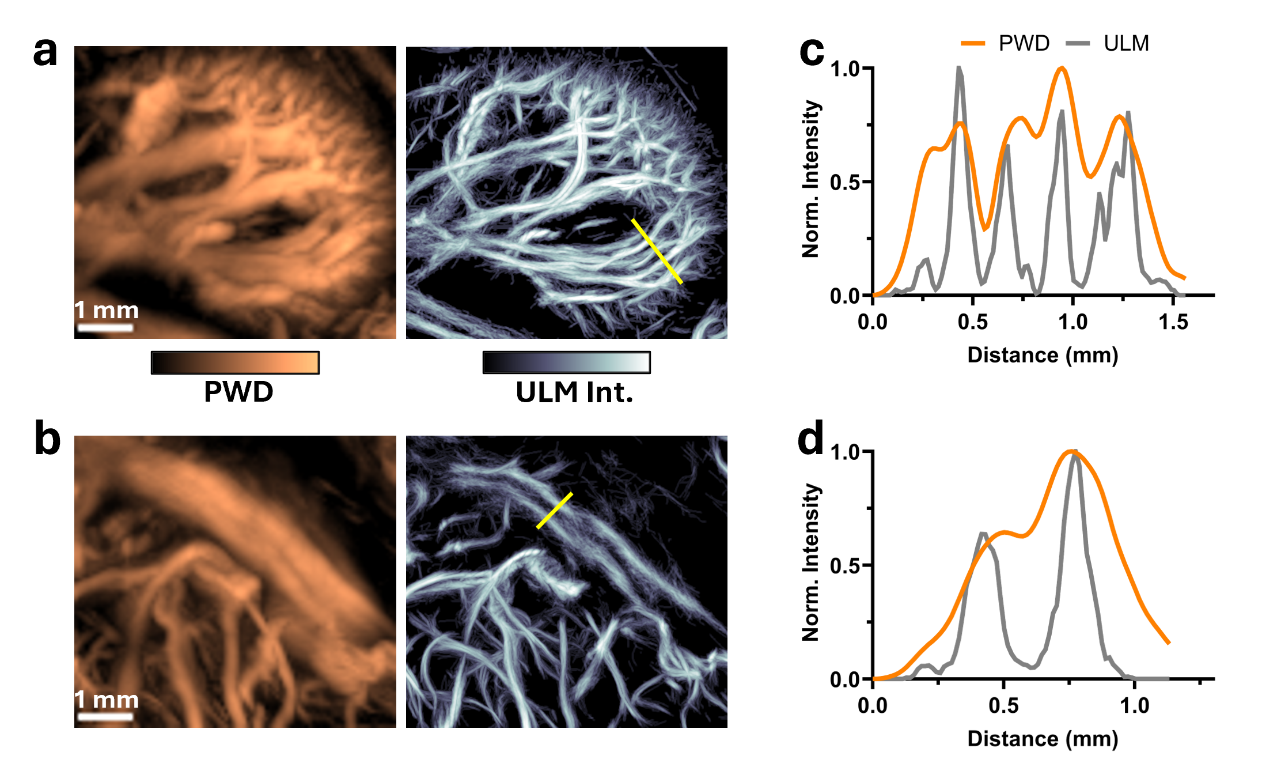

**Supplementary Figure 8. Comparison of vascular delineation by PWD and ULM.** Matched PWD and ULM intensity images of **(a)** the left kidney and **(b)** the iliac vessels. **(c, d)** Normalized intensity profiles extracted along the yellow lines in panels **(a)** and **(b)**, respectively. Compared with PWD, ULM produced sharper vessel profiles and improved separation of adjacent vascular features, allowing more microvessels to be distinguished.

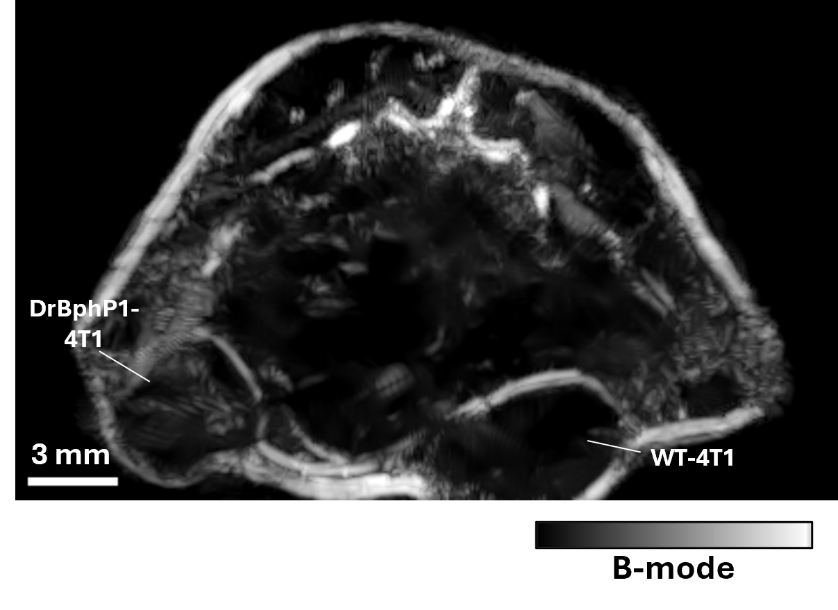

**Supplementary Figure 9. B-mode visualization of the bilateral tumor xenografts***.* Cross-sectional B-mode image acquired at the elevational position indicated by the white dashed line in **Fig. 4d**. The boundaries of the DrBphP1-4T1 tumor and the contralateral WT-4T1 tumor are delineated by acoustic-scattering contrast.

**Supplementary Video 1. Whole-body volumetric photoacoustic imaging at 870 nm.** Three-dimensional rendering of the mouse torso acquired in vivo using PACT at 870 nm. Elevational scanning enables neck-to-tail visualization of blood-rich structures throughout the thoracic and abdominal regions.

**Supplementary Video 2. Whole-body volumetric B-mode ultrasound imaging.** Three-dimensional US B-mode rendering of the mouse torso acquired in vivo. The volumetric dataset delineates the external body boundary and internal anatomical structures.

**Supplementary Video 3. Whole-body volumetric power Doppler imaging.** Three-dimensional rendering of MB-enhanced PWD imaging of the mouse torso acquired in vivo. The volume visualizes perfused vasculature from the thoracic cavity through the abdominal and lower-abdominal regions.

**Supplementary Video 4. Whole-body sliding-depth photoacoustic imaging at 870 nm.** Successive transverse PACT MAPs of the mouse torso acquired in vivo at 870 nm using a sliding depth window of 1 mm. The video progresses from the neck to the lower abdomen and visualizes blood-rich structures throughout the imaging volume.

**Supplementary Video 5. Whole-body sliding-depth B-mode ultrasound imaging.** Successive transverse B-mode MAPs of the mouse torso acquired in vivo using a sliding depth window of 1 mm. The video progresses from the neck to the lower abdomen and delineates the external body boundary and internal anatomical structures based on acoustic-scattering contrast.

**Supplementary Video 6. Whole-body sliding-depth power Doppler imaging.** Successive transverse MB-enhanced PWD MAPs of the mouse torso acquired in vivo using a sliding depth window of 5 mm. The video progresses from the neck to the lower abdomen and visualizes perfused vasculature across the thoracic and abdominal regions.

**Supplementary Video 7. Sliding-depth power Doppler imaging overlaid with differential photoacoustic signal.** Successive transverse MAPs of the bilateral tumor region acquired using a sliding depth window of 3 mm. MB-enhanced PWD imaging visualizes the surrounding perfused vasculature, while the overlaid differential photoacoustic signal highlights the DrBphP1-expressing 4T1 tumor relative to the contralateral wild-type 4T1 tumor.

**Supplementary Video 8. Sliding-depth B-mode ultrasound imaging of tumors.** Successive transverse B-mode MAPs of the bilateral tumor region acquired using a sliding depth window of 1 mm. Acoustic-scattering contrast delineates the DrBphP1-expressing 4T1 tumor and the contralateral wild-type 4T1 tumor, together with surrounding anatomical structures.

**Supplementary Video 9. Sliding-depth photoacoustic imaging of tumors at 870 nm.** Successive transverse PACT MAPs of the bilateral tumor region acquired at 870 nm using a sliding depth window of 1 mm. The video visualizes hemoglobin-associated PA contrast within and surrounding the tumors across successive depths.
